# NDNF+ interneurons have a privileged role in regulating cortical excitability

**DOI:** 10.1101/2025.11.27.690945

**Authors:** Amy Richardson, Marion S. Mercier, Yoshiteru Shimoda, Robert T. Graham, Marco Leite, Alejandro Garcia, Maud Muller, Suraya A. Bond, Maria Grozdanova, Qimin Wu, Andreas Lieb, Dimitri M. Kullmann, Vincent Magloire

## Abstract

Cortical processing depends critically on the precise modulation of excitatory activity by a diverse population of GABAergic interneurons. Among these, neuron-derived neurotrophic factor-expressing (NDNF+) interneurons have been proposed to be "master regulators" of cortical microcircuits. While their role in coordinating physiological cortical activity is beginning to be understood, their contribution to restraining hyperexcitability during epileptiform activity remains unexplored. To address this, we employed calcium imaging, optogenetics, and chemogenetics in an NDNF-Cre mouse line to investigate how NDNF+ interneurons influence cortical hyperactivity and spontaneous seizures. Our results demonstrate that NDNF+ interneurons are actively recruited during interictal spikes and focal seizures, although more slowly than parvalbumin-positive interneurons. Optogenetic hyperpolarization of NDNF+ interneurons exacerbates epileptic discharges, whereas their depolarization suppresses focal seizures, even when the optogenetic stimulus is delayed by several seconds from seizure onset. This effect is largely mediated by GABAB receptor signalling. Additionally, chemogenetic depolarization of NDNF+ interneurons has robust antiepileptic effects both ex vivo and in vivo. Collectively, these findings establish NDNF+ neurons as key regulators of cortical excitability and identify them as promising cellular targets for the development of anti-epileptic therapies.

## Introduction

Cortical computations depend critically on the GABAergic inhibitory system, which governs the flow of information through the cortex with high spatiotemporal precision. Somatic inhibition exerts a strong control over pyramidal neuron output by modulating action potential generation and timing, while dendritic inhibition regulates the integration of synaptic inputs across the dendritic arbour^1^.

Over 20 subtypes of GABAergic interneurons have been identified, each exhibiting distinct electrophysiological, morphological, genetic, connectivity, and GABA signalling profiles^2–4^. These subtypes have been broadly classified into three groups based on the expression of molecular markers, with parvalbumin-positive (PV+), somatostatin-positive (SOM+), and vaso-active intestinal polypeptide-positive (VIP+) interneurons accounting for 80-90% of all interneurons^1^. PV+ fast-spiking interneurons, comprising approximately 40% of cortical interneurons, target the peri-somatic region of pyramidal neurons. They mediate fast, transient feedforward and feedback inhibition critical for the generation of gamma oscillations and the temporal segregation of neuronal assemblies^1,5–8^. SOM+ interneurons, accounting for roughly 30% of cortical GABAergic cells, inhibit the distal dendrites of pyramidal neurons. They deliver slower-onset, sustained inhibition and are primarily involved in feedback inhibition, modulating dendritic excitability across cortical layers and cell types^1,9,10^. As for VIP+ interneurons, they primarily disinhibit cortical pyramidal neurons by inhibiting SOM+ interneurons^1,11–13^. This disinhibition creates spatio-temporal windows that facilitate pyramidal neuron firing, thereby broadly influencing cortical network excitability^1,14–16^.

A fourth group of GABAergic interneurons has recently been genetically defined, mainly comprising neurogliaform (NGF) neurons. This class is distinguished by the expression of neuron-derived neurotrophic factor (NDNF) or inhibitor of DNA binding 2 (Id2) markers, depending on their location within the cortical column^17–20^. NGF neurons provide long-range feedforward inhibition and exhibit unusual synaptic properties, including a high density of GABA release sites along their profuse axons and a relatively wide synaptic cleft^21^. These features enable them to signal at least in part via "volume transmission"^21^. Characterized by the release of large GABA clouds into the extracellular space, this mechanism allows a single NGF cell action potential to induce a powerful, long-lasting inhibitory response in pyramidal neurons via both long-lasting GABAA (GABAA_slow_) and GABAB signalling^19,21,22^. Beyond these signalling properties, NGF neurons exhibit a high connection probability across cell types and cortical layers^9,19,23^ and form a syncytium through gap-junction coupling, not only among themselves but also with other interneurons^24,25^. This enables them to act as "master regulators" of the cortical microcircuit^9,26^, capable of modulating the excitability of entire cortical columns.

Cortical inhibition plays a pivotal role in regulating cortical excitability under physiological conditions through the coordinated action of these four interneuron classes. However, the contribution of each population to restraining excitation during excessive or paroxysmal activity, such as epileptiform discharges, is less clear. While it is established that a failure of inhibition underlies seizure generation and propagation into new cortical territories^27–32^, which interneuron subtypes prevent (or promote) hyperexcitability, and by which mechanisms, is still incompletely understood. The identification of distinct molecular markers, combined with advances in opto- and chemogenetic tools as well as cellular imaging, has revealed that the inhibitory control exerted by different interneuron subtypes varies significantly during epileptic activity.

For instance, PV+ cells, while being recruited very early during epileptic discharges^28,29,33–36^, can suppress or promote cortical excitation depending on when and where they are activated relative to the hyperexcited pyramidal network ^35,37–48^. In contrast, SOM+ interneurons are activated later during seizure activity^29,34–36^ and generally decrease network excitability when optogenetically stimulated, although this effect can wane^35,37,38,45,47–49^. Few studies have examined the role of VIP+ interneurons in restraining excitation during epileptic activity, but in vivo optogenetic inhibition of VIP+ cells has been shown to increase seizure threshold^35^ and silencing their synaptic activity has anti-seizure effect^50^, suggesting an overall reduction in cortical excitability. Conversely, chemogenetic activation of VIP+ interneurons in an ex vivo model of epileptiform discharges produced no detectable effect^49^. The role of NGF neurons in this context remains largely unexplored. These neurons express neuropeptide Y (NPY),having anti-epileptic properties^51,52^, and exhibit atypical persistent barrage firing in the absence of synaptic inputs, a mechanism thought to counteract over-excitation^26,53^. However, the lack of identified molecular markers specific to NGF interneurons has until recently^17–19^ hindered investigation of their roles in states of circuit hyperactivity such as epilepsy.

Here, we leverage a recently generated NDNF-Cre mouse line, combined with calcium imaging, optogenetics, and chemogenetics, to investigate the role of NDNF+ NGF interneurons in controlling hyperexcitability and spontaneous seizures in multiple ex vivo and in vivo epilepsy models. We show that NDNF+ interneurons are actively recruited during states of cortical hyperactivity including interictal spikes and focal seizures, albeit more slowly than PV+ cells. Optogenetic hyperpolarization of NDNF+ interneurons promotes epileptic discharges, whilst optogenetic depolarization has the opposite effect, even when delayed several seconds after seizure onset, an effect that contrasts sharply with the narrow temporal window of PV+ and SOM+ inhibition^37^. Collectively, these results strongly implicate NDNF+ NGF neurons as privileged regulators of cortical excitability even in pathological states. Finally, chemogenetic excitation of NDNF+ interneurons had an antiepileptic effect both in ex vivo cortical and hippocampal slice preparations, and in an in vivo temporal lobe epilepsy model. The results identify NDNF+ interneurons as a potential cellular target for novel anti-epileptic therapies.

## Materials and Methods

### Animals and Ethical approval

All experimental procedures were carried out in accordance with the Animals (Scientific procedures) Act 1896. Mice were housed under a 12h:12h light-dark cycle and had access to food and water *ad libitum*.

Heterozygous and homozygous NDNF-Cre mice (B6.Cg-Ndnf^tm1.1(folA/cre)Hze^/J; JAX 028536) were locally injected with adeno-associated viral vectors carrying floxed genes of interest. Heterozygous PV::Cre (B6;129P2-Pvalb^(tm1(cre)Arbr)^/J mice; JAX 008069) and SOM::Cre (B6N.Cg-Sst^tm2.1(cre)Zjh/J^; JAX 013044) mice were either crossed with Ai32 mice (B6;129S-Gt(ROSA)26Sor^tm32(CAG-COP4*H134R/EYFP)Hze/J^; JAX 012569) for conditional expression of EYFP-tagged ChR2 in PV+ and SOM+ target interneurons, or locally injected with adeno-associated viral vectors ^37^.

### Plasmid construction

#### AAV-mDlx-FLEX-hM3D(Gq)-mCherry

The pAAV-mDlx-FLEX-hM3Dq-mCherry transfer plasmid was created by replacing the hsyn promoter in pAAV-hsyn-DIO-hM3D(Gq)-mCherry (Addgene #44361) with an mDlx promoter from pAAV-mDlx-GCaMP6F-Fishell-2 (Addgene #83899). This was achieved by inserting flanking MluI and SalI restriction sites on either side of the mDlx promoter using the primers: Fwd – GATACGCGTctatacactcacagtggtttgg (MluI) and Rev - GATGTCGACctgtggagagaaaggc (SalI). Standard restriction-based cloning was used to insert the mDlx promoter fragment into the pAAV-FLEX-hM3Dq-mCherry backbone.

#### pAAV-mDlx-FLEX-mCherry

The pAAV-mDlx-FLEX-mCherry control plasmid was created by removing hM3Dq transgene from the pAAV-mDlx-DIO-hM3Dq-mCherry plasmid using Q5 site-directed mutagenesis (NEB E0554S) with the primers Fwd – GGTGGCGCTAGCATAACTT and Rev – GCCACCATGGTGAGCAAG.

#### pAAV-mDlx-FLEX-Chronos-GFP and pAAV-mDlx-FLEX-ArchT-GFP

mDlx promoters replaced EF1α and CAG in the source plasmids pAAV-EF1α-FLEX-Chronos-GFP (Addgene #62725) and pAAV-CAG-FLEX-ArchT-GFP (Addgene #28307), respectively. The mDlx promoter placed into the Chronos-GFP backbone was taken from the source plasmid (Addgene #83899) using the following promoters to insert PacI and EcoRI restriction sites; Fwd-GATTTAATTAActatacactcacagtggtttgg and Rev – GATCGAATTCcctgtggagagaaaggc. Standard restriction-based cloning was used to insert the mDlx promoter fragment into the pAAV-FLEX-Chronos-GFP backbone. The mDlx promoter was amplified from the pAAV-mDlx-FLEX-mCherry plasmid and inserted into the pAAV-FLEX-ArchT-GFP backbone using a NEB HiFi DNA assembly protocol. The primers used to amplify the mDlx promoter sequence were Fwd-atgttcccatagtaacgccaGTCTATACACTCACAGTGG; Rev-atggggatccaattctttgccTGTGGAGAGAAAGGCAAAG. Those used to amplify the backbone sequence were Fwd-GCAAAGAATTGGATCCCC; Rev – TGGCGTTACTATGGGAAC.

### Viral Vectors

All plasmids listed above were fully sequenced before AAV production by VectorBuilder. The complete list of AAVs as well as their specific use for the present study are given in **Suppl. Table 1**.

### Surgical procedures

#### Viral vector injections

Mice of both sexes (2-6 months old) were anaesthetised with 5% isoflurane, and transferred to a stereotaxic frame (David Kopf Instruments Ltd, USA) after shaving the fur on the head. Animals were injected with buprenorphine (0.1 mg/kg, s.c.) and Metacam (2 mg/kg, s.c.) for pain relief. Core temperature was maintained using a thermoblanket, and anaesthesia was maintained with 1.5-2.5 % isoflurane. After confirming loss of the pedal reflex, an incision was made to the scalp and burr holes were drilled at different locations depending on the type of experiment (**Suppl. Table 1**). For cortical injections, the tip of a 34-gauge bevelled needle (12°, 25 mm length; Esslab Ltd., UK) was inserted at an angle of 30° to the pial surface, with the bevel facing upward to maximise the viral load to the superficial cortical layers. Coordinates, volume, AAV pseudotypes and titres used for each experiment are given in **Suppl. Table 1**. Viral constructs were injected at 100 nL / min using a micro-injector (WPI) connected to a Hamilton syringe (5 μL RN; Hamilton Company; #7647-01). The needle was left in situ for 5 minutes after injection before withdrawing it. The scalp was then sutured, and animals were injected with sterile saline (0.2-0.3 mL, s.c.), and allowed to recover in a heated chamber. Animals were allowed to recover for at least a week before experiments or further surgeries.

#### In vivo calcium imaging

Three weeks after viral injection in the upper layer of the visual cortex, NDNF or PV-Cre mice underwent a second surgery to affix a headplate (Model 5, Neurotar Ltd.) as described^54^. In addition to buprenorphine and Metacam, mice were given dexamethasone (2 mg/kg, i.m.). A cranial window was performed above the visual cortex (**Suppl. Table 1**). The craniotomy was initially outlined with a 2 mm biopsy punch (Kain Medical) followed by skull thinning using a right-angled dental drill (NSK) with a diamond burr (Diatech). Sterilised 2 mm silicon disks (Sylgard 84; 0.31 mm thickness from ref. ^55^) were secured to a 3 mm glass coverslip with a UV curing optical adhesive (Norland 68) and placed inside the craniotomy to replace the arachnoid space and avoid formation of air bubbles. Coverslips were glued to the skull using cyanoacrylate adhesive (Vetbond).

Posterior to the cranial window, a small burr hole was drilled and a stainless-steel wire was implanted at the cortical surface to record the electrocorticogram (ECoG). The chemoconvulsant was delivered via the same burr hole on the day of experiment. A reference electrode was placed on the contralateral frontal lobe. The headplate and electrode connector were affixed to the skull with opaque dental cement (Super-Bond, Sun Medical) and the entire chamber was covered with an opaque silicon-elastomer sealant (Kwik-Cast, WPI Europe) to protect the cranial window. Animals were allowed to recover for 5 days before habituating to the head-fixed system for increasing periods (10 to 40 minutes) over at least 7 days.

#### In vivo optogenetic and chemogenetic experiments

During the same surgery to inject AAVs optical adhesive (Norland 68) was applied to the skull above the cortex of interest in preparation for a future cranial window. A headplate (model 5 or 13 Neurotar, Ltd) was affixed to the skull using opaque dental cement (Super-bond, Sun Medical) and the site was covered with a layer of opaque silicone elastomer (Kwik-cast, WPI Europe). Mice were injected with saline (0.2-0.3 mL, s.c.) and allowed to recover for at least five days before habituation to the head-fixed system. During habituation, water or 30% sucrose was offered to the mice via 1 mL syringe with a blunt end needle.

Three to four weeks later, on the day of the experiment, mice underwent a second surgery to perform a cranial window above the cortex of interest. Mice were anaesthetised and transferred to a stereotaxic frame as described above, and injected with Metacam (2 mg/kg, s.c.) and dexamethasone (2 mg/kg, i.m.). The Kwik-cast and optical adhesive were removed from the skull and a 2 mm craniotomy was then performed as detailed above.

For the optogenetic experiments, the cranial window was not covered but was irrigated with warm sterile saline and covered with silicone elastomer (Kwik-cast). For chemogenetic experiments, sterilised 2 mm silicon disks were placed inside the craniotomy as above. In all cases, animals were injected with 0.2-0.3 mL saline and allowed to recover for several hours before starting recordings.

#### In vivo intrahippocampal kainic acid, viral injection and LFP electrode implantation

Mice of both sexes (3-4 months old) underwent two surgeries, the first to induce status epilepticus (and consequently establish chronic epilepsy), and the second to inject AAVs and implant depth electrodes and a subcutaneous wireless transmitter to record LFP signals in the hippocampus. Mice were anaesthetised with isoflurane (5 %), and transferred to a stereotaxic frame, and injected with buprenorphine and Metacam as above. They were also injected with bupivacaine (0.02 mL of 0.25 % Marcain diluted 1:1 in sterile saline, s.c.) to the scalp as an incisional block. A burr hole was drilled above the dorsal hippocampus (coordinates from bregma, AP: -2.8 mm; ML: 2 mm; DV: -2 mm) and kainic acid (synthetic, Tocris; 7 mM, 140 nL in males and 7 mM, 80 nL females) was injected into the dentate gyrus at a rate of 200 nL/min (Hamilton syringe, Hamilton Company; 33-gauge flat needle, Esslab Ltd, UK). The needle was retracted two minutes after the end of the injection. The scalp was sutured and mice were administered 0.3 mL saline with glucose (4% w/v, s.c.). Mice were then placed in a heated recovery chamber for 15 minutes before being returned to their home cage, where they were monitored for 2 hours to assess the severity of status epilepticus. Either 2 hours after kainic acid injection or after occurrence of three Racine Stage 5 seizures, whichever occurred sooner, mice were given diazepam (10 mg/kg, i.p.) to terminate status epilepticus (SE). Racine stage 5 seizures were defined as loss of righting reflex following rearing and clonus and/or wild jumping^56^.

Two weeks later, mice that had exhibited Racine stage 5 seizures were injected with AAV9-mDlx-FLEX-mCherry or AAV9-mDlx-FLEX-hM3Dq-mCherry and implanted with a wireless transmitter (Open Source Instruments, Single channel, 256 Hz either A3028C-AA-B45-B or A3048S2-AA-C45-D). Briefly, mice were anaesthetised with 5% isoflurane, transferred to a stereotaxic frame and injected with buprenorphine and Metacam as above. AAVs were injected bilaterally in the stratum lacunosum moleculare (SLM) of the hippocampus as above. Following viral injection, a small incision was made to the back of the mouse and a wireless transmitter was placed subcutaneously. The leads were tunnelled under the skin through to the skull and a depth electrode was attached to the recording electrode via a metal crimp (Open Source Instruments, SDE-X). A burr hole was drilled (coordinates AP: - 3.1 mm; ML: 3.12 mm) and a Teflon-coated stainless steel wire depth electrode was lowered to a DV of -1.5 mm below the pia, 1 mm above the viral injection site in the ventral hippocampus. A reference electrode was placed on contralateral frontal cortex. The animal was sutured and the electrodes sealed in place with dental cement (Simplex Rapid, Kemdent). Animals were given 0.3 mL saline and placed in a heated recovery chamber, before returning to their home cage where they were allowed to recover for 2-3 weeks before recordings.

### Calcium imaging experiments

#### Acquisition

Imaging data was recorded using an Olympus FV1200MPE multi-photon laser scanning microscope with a 25x immersion objective (XLPLN25XWMP2, Olympus) while head-fixed on a floating platform (Neurotar Ltd.) Mice were habituated to the recording setup for a week before recordings. GCaMP6f and mCherry were excited at 910 nm and 800 nm respectively using a Ti-Sapphire Chameleon Ultra Pulsed Laser (Coherent). Emission fluorescence was collected using two photomultipliers (PMTs) and filters (PMT 1: 515–560 nm; PMT2: 590–650 nm). Imaging was performed using 254 x 254μm frame scans (128x128 or 256x256 pixels) at a frame rate of 15 or 8Hz respectively. Resolution and power were kept constant within each experiment and each image acquisition lasted 60-90s.

To image calcium-dependent fluorescence of NDNF+ and PV+ cells during seizures, mice were anaesthetised with 1.5 % isoflurane and transferred to a stereotaxic frame for injection of the chemoconvulsant. Kwik-cast was removed from the headpiece and pilocarpine (3.5 M, 150 nL, Tocris) was injected via the burr hole located next to the ECoG wire into the visual cortex at a depth of 0.5-0.6 mm and a speed of 100 nL/min using a Hamilton syringe needle (33-gauge, flat end). Mice were then transferred to the head-fixed imaging setup for the recording session. Three to four imaging windows were recorded in each animal approximately 1.5 mm from the chemoconvulsant injection site and ECoG recording wire.

The coordinates for each imaging site were recorded from the x,y coordinates a motorised platform (Scientifica) holding the head-mounting arm. The concomitant ECoG signal was acquired using a Multiclamp 700B amplifier (Molecular Devices), digitised at 10 KHz, bandpass filtered between 0.1 Hz – 1 KHz, and recorded using WinEDR (Strathclyde University, UK). The signal was time-stamped with imaging data by recording a TTL signal from the scanning head. ECoG recordings were initiated at the time of chemoconvulsant injection so that the latency to seizures could be monitored. Experiments typically lasted for 40-50 minutes and animals were monitored closely for welfare. Mice were offered water and 30% sucrose solution as above. If electrographic seizure activity was observed at the end of the recording session, diazepam (10 mg/kg, i.p.) was administered before returning the mouse to its home cage. In animals where pilocarpine failed to initiate seizures, a second recording session was performed a few days later.

For in vivo calcium imaging combined with chemogenetic activation (**Suppl. Fig. 11**), at least three hours after surgery, mice were transferred to the head-fixed imaging system and the Kwik-cast was removed from the headpiece. Two to three imaging windows were recorded sequentially in each animal. A baseline was acquired by performing 5 acquisition periods (180 s each) taken 5 minutes apart. The animal then received either DMSO in saline (1 %, i.p.), deschlorclozapine (DCZ, 10 μg/kg in 1% DMSO) or clozapine (CLZ, 100 μg/kg in 1% DMSO). Image acquisitions resumed 10 minutes after the injection and were repeated every 5 minutes for a further 40 minutes. At the end of the recording session, the silicone elastomer was replaced on the headpiece and the mouse was returned to its home cage.

The experiment was repeated over the course of three consecutive days and each day the mice received a different drug or DMSO in saline control. The order of the drug injected for each animal was randomised and the experimenter was blinded to both the virus and the drug injected.

#### Analysis

Image analysis was done using in-house custom MATLAB scripts. Images from each recording session went through a series of pre-processing steps including frame registration using 2D cross-correlation to correct for small X-Y movements that inevitably occur during seizures and flat-field correction to correct any vignetting. Top-hat filtering using a disk structuring element was done to enhance brightness in objects that had the approximate size and shape of neuronal somata. Regions of interest (cell bodies) were then selected for analysis only if they were mCherry positive. Fluorescence signals were extracted from each pixel within the ROI, ignoring those at the soma boundary, to decrease potential contamination from surrounding neuropil. A local background calculated from the area surrounding a cell soma was subtracted from the cell signal and ΔF/F was calculated using the median cell signal.

Each fluorescence point was aligned to the centre of the TTL pulse in the electrophysiological recordings. Seizures were analysed if they lasted > 2 s and spikes were larger than 5 standard deviations (SD) above the noise level. Calcium transients were analysed if they were > 2x SD above background noise. An exponential curve was fitted to the calcium transient to calculate ‘time to plateau’.

### In vivo optogenetic experiments

#### Acquisition

Two to three hours after craniotomy surgery, pilocarpine was locally injected (3.5 M, 150 nL, 100 nL / min) with a Hamilton syringe and a 34-gauge flat needle (Esslab) at the centre of the craniotomy at a depth of 0.5 mm under isoflurane anaesthesia (1.5-2 %). The LFP recording system was activated at the start of the injection to create a timestamp. Three minutes after the injection the needle was retracted and the animal placed in the head-fixed system without further anaesthesia. Typically, within 5 minutes, the animal would be fully awake. In the meantime, an LFP recording electrode (borosilicate glass pipettes 1.5 mm OD, 0.86 mm ID Warner Instruments, G150F-4 with a resistance of 2-3 MΩ) filled with saline was positioned at the centre of the craniotomy above the pilocarpine injection site and lowered to 0.3 mm below the pia, and a bare ended optic fibre (ø: 200 µm, NA: 0.22, Multimode 190–1200 nm, Thorlabs Inc., UK) was positioned 1.5-2 mm above the pia (**Suppl. Fig. 1A**). The entire craniotomy and chamber created by the headplate was filled with warm saline and a silver-wire coated with chloride was added to the bath for referencing.

The LFP signal was acquired using a Multiclamp 700B amplifier, filtered between 1 and 300 Hz with a gain of 50 (mV/mV) and digitized at 1 kHz using a NI board and recorded with WinEDR. The LFP signal was passed to a field-programmable gate array (FPGA) CRIO-9076 integrated controller (National Instruments, USA) which triggered the optogenetic laser system in real-time upon seizure detection. A laser source either at 470 or at 570 nm (CNI laser) was used to activate a Chronos or ArchT construct respectively. The power at the tip of the fibre was between 10 and 15 mW and kept constant throughout the experiment.

Ambient light in the room was kept constant and fairly bright to occlude indirect stimulation of the retina. Indeed, preliminary results in dim light conditions suggested that laser stimulation could occasionally entrain activity in the visual cortex in the absence of opsin expression (**Suppl. Fig. 1B**).

Optogenetic stimulation started when the LFP signal was exhibiting interictal spikes (about 20-30 minutes post-injection as previously reported^27,37^). Each stimulation lasted 10 s and was interspersed by >30 s non-stimulation periods. For ArchT experiments, continuous illumination was applied while various frequencies of pulsed stimulation were used for the Chronos experiment. For Chronos activation, 2, 4, 8, 10, 20, 50, and 100 Hz pulses (duty cycle 50:50) were applied pseudo-randomly throughout the experiment. For tests of interictal spike optogenetic manipulation, laser stimulation was applied in open-loop. To test the effects of optogenetic manipulations on focal seizures, laser stimulation was applied in closed-loop. Briefly, the sentinel spike at the start of seizure onset was detected via a custom Labview threshold-crossing script run on the FPGA, allowing the electrographic seizure to be detected within ∼100 µs. Upon detection, the laser stimulation (either continuous or pulsed) was activated for 10 s with or without a delay. Finally, during closed-loop stimulation, optogenetic stimulation (laser on) was alternated with sham stimulation (detection of the event without laser on).

At the end of the experiment, animals were culled and transcardially perfused with PFA. The fixed brains were then sliced to 100 μm thickness and analysed to verify expression of the optogenetic actuator and quantify its spread (typically between 2 and 3 mm along the postero-anterior axis, **Suppl. Fig. 1C**). Ectopic expression of viral constructs in pyramidal neurons was estimated by counting the number of fluorescent pyramidal shaped cells within all 100 μm slices containing the viral construct.

#### Analysis

All the data were analysed using in-house custom Python scripts. After an initial correction for artefacts, LFP traces were down-sampled to 100 Hz, and filtered using a Butterworth bandpass filter from 1 to 30 Hz. The onset of interictal discharges was determined by using a detection threshold on the second derivative of the LFP recording. The threshold corresponded to a minimum of 5 SDs above baseline noise. The seizure was also detected using detection threshold on the second derivative of the LFP recording with a minimum of 3 SDs above baseline noise. Durations and delay between seizure onset and light stimulation was then inferred.

### Chemogenetic activation of NDNF+ cells in chronic kainate epilepsy model

#### Acquisition

Two weeks after viral injection and LFP transmitter implantation, transmitters were switched on to record at least 1 hour of baseline. Mice were then injected with either DCZ (10 μg/kg in 0.1% DMSO, Tocris), i.p. or DMSO (0.1 %, Sigma). LFP was recorded for at least 60 minutes post-injection before switching transmitters off. Two days later, the transmitter was turned on for a baseline recording of 1 hour before intraperitoneal injection of either DMSO or DCZ (so that each mouse received an injection of both compounds across the two days). The experimenter was blinded to both the virus and drug injected.

#### Analysis

One hour of LFP recording pre- and post-injection was analysed. LFP recordings were annotated using in-house software (PyECoG2, https://www.pyecog.com/) to classify electrographic activity into 3 types of events – seizures, interictal activity and baseline.

Seizures were classified as multiple spiking events that were 2X the standard deviation of the noise and lasted at least 10 s. Two events that occurred within 5 s of each other were counted as one seizure. Baseline was 30 s of recording that contained 6 spikes or fewer. Interictal activity was defined as multiple spike events that lasted < 10s. Periods of discontinuous LFP signal were discarded.

### *Ex vivo* electrophysiology

#### Brain slicing

Male and female NDNF-Cre mice aged 2-4 months injected with AAVs (**Suppl. Table 1**) were briefly anaesthetised and given a terminal dose of intraperitoneal sodium pentobarbital (500 mg/kg). Following cardiac perfusion with ice cold sucrose solution, the brain was removed. Sucrose solution contained (in mM): Sucrose (75), NaCl (87), Glucose (24), NaHCO3 (26), KCl (2.5), MgCl (7) NaH2PO4 (1.25) and CaCl2 (0.5), oxygenated with carbogen (95% O2 / 5% CO2).

For cortical slices, sagittal slices were made at 300-400 μm thickness using a vibrating microtome (Leica VT1200S, Leica). For hippocampal slices, 400 μm horizontal sections were cut as described^57^. All slicing was performed in ice-cold slicing solution bubbled with 95% O_2_ / 5% CO_2_. Slices were allowed to recover for 30 minutes in a slicing solution warmed to 35 °C and then allowed to rest at room temperature for 30 minutes prior to experiments.

Slices used for *ex vivo* seizure models were transferred to an interface chamber filled with storage solution containing: NaCl (119), NaHCO_3_ (26), Glucose (10.92), KCl (2.5), NaH_2_PO_4_ (0.93), CaCl_2_ (2) and MgCl_2_ (1). The chamber was provided with humidified carbogen.

#### Local field potentials and ex vivo seizure models

##### Acquisition

Slices were suspended on a net within a recording chamber on the stage of an upright microscope. This allowed the recording solution to perfuse both sides of the slice, improving the reliability of initiating seizure-like events. Slices were continuously perfused with a low Mg^2+^ recording artificial cerebro-spinal fluid (aCSF) containing (in mM): NaCl (119), NaHCO_3_ (26), Glucose (22), KCl (3.5) and CaCl_2_ (2.5) at 10 mL / min, heated to 35 °C. For experiments using cortical slices 0.05 mM MgCl_2_ was included in the recording aCSF, while for those using hippocampal slices 0.15 mM MgCl_2_ was added. Slices were visualised using an upright microscope (Olympus BX51WI) equipped with differential interference contrast illumination, a water immersion 20x objective (Olympus XLUMPlan FLN, 1.00 NA) and a CCD camera (Ikegami, ICD-47E). Prior to each slice experiment, mCherry fluorescence expression was checked using a 590 nm LED (Thorlabs M590L2) with a mCherry filter cube (Semrock, mCherry-40LP-A-OMF).

LFP recording was performed either from layer 5 of the visual cortex, in a region with mCherry fluorescence in layer 1, or from layer 5 of the entorhinal cortex where mCherry fluorescence was present in the SLM of the hippocampus. LFP was recorded using borosilicate glass pipettes (1.5 mm OD, 0.86 mm ID Warner Instruments, G150F-4), with a resistance of 2-3 MΩ, filled with recording aCSF. LFP signals were acquired with a Multiclamp 700B amplifier connected to a National Instruments board (NI USB-6341) and a computer running WinWCP (University of Strathclyde). Signals were low pass filtered at 1 KHz and digitized at 10 KHz.

A tungsten concentric bipolar electrode (World Precision Instruments, TM33CCINS) was used to electrically stimulate layer 5 of the visual cortex or layer 2/3 of the entorhinal cortex to generate seizure-like events (SLEs) from a defined focal region. Electrical stimulation trains (100 Hz for 1 s, 0.1 ms step) of increasing current intensity (20 μA-200 μA) were applied in 20 μA increments via constant current stimulator (Digitimer Ltd, DS3 689) until a SLE was initiated. A SLE was defined as a minimum of 4 consecutive short discharges. The current required to elicit an SLE was defined as the threshold and 110% of this threshold was then used subsequently to elicit SLEs reliably. Slices were allowed to rest for 10 minutes following the first SLE. Test SLEs were elicited every 5 minutes with repeated electrical stimulation trains for a total of 12 cycles. Recording aCSF containing the hM3Dq agonist clozapine-N-Oxide (CNO, 10 μM, Cambridge Bioscience CAY25780) was applied immediately following the 6^th^ stimulation.

##### Analysis

Average parameters from SLEs generated by stimulations 1-6 formed the ‘baseline response’ and averaged parameters from SLEs at stimulations 10-12 formed the ‘treatment response’. To allow time for diffusion of CNO into the slice, stimulations 7-9 were not analysed. SLEs were analysed using in-house custom Python scripts. Briefly, recordings were low pass filtered at 40 Hz and spike detection was performed on 2-minute portions of the recording immediately following an electrical stimulation. SLE detection consisted of counting spike events 5X larger than the SD of the baseline noise. The experimenter was blinded to the virus injected throughout the recording and analysis period.

#### Whole-cell patch clamp recordings

##### Acquisition

Following a thirty-minute recovery period at room temperature, slices were transferred to a submerged recording chamber and continuously perfused with aCSF containing (in mM): NaCl (125), NaHCO_3_ (25), D-glucose (25), KCl (2.5), NaHPO_4_ (1.25), CaCl_2_ (2) and MgCl_2_ (1), heated to 35 °C, at 2 mL/min. Slices were held in place using a platinum harp with nylon threads and visualised using an upright microscope as above. mCherry or GFP fluorescence was used to identify NDNF+ interneurons expressing either chemogenetic, or optogenetic actuators as well as control reporters. Borosilicate glass pipettes (1.5 mm OD, 0.86 mm ID Warner Instruments, G150F-4) with a resistance of 2.5-3.5 MΩ and filled with a solution containing (in mM): K-gluconate (122), KCl (6), HEPES (10), Na-phosphocreatine (10), Mg-ATP (4), Na-GTP (0.4), EGTA (0.2) and MgCl2 (1) were used to record intrinsic and active electrical properties of NDNF+ interneurons. Current clamp recordings were achieved using a Multiclamp 700B amplifier (Molecular Devices) connected to a National Instruments board (NI USB-6341) and a computer running WinWCP (University of Strathclyde) or an in-house NI acquisition software. Signals were filtered at 10 kHz and digitized at 50 kHz. Cells that had a series resistance >25 MΩ or a resting membrane potential > -50mV were not included.

##### For optogenetic depolarisation and hyperpolarisation construct validation

For the characterisation of optogenetic activation of NDNF+ cells, trains of 1 ms-long light stimuli, with 5 pulses at 1 Hz, 10 pulses at 2 Hz, 20 pulses at 5 Hz, 20 pulses at 10 Hz, 20 pulses at 20 Hz, 20 pulses at 50 Hz, 20 pulses at 80 Hz and 20 pulses at 100 Hz, were delivered three times consecutively, with inter-train intervals of 15 s. This was done while maintaining the resting potential at -70 mV in current-clamp mode. For the characterisation of inhibitory post-synaptic potentials (IPSPs) in pyramidal neurons elicited by NDNF+ cell photo-activation, the following protocol was looped twice with a 60 s long inter-stimulus interval: 15 pulses at 5 Hz, 30 pulses at 10 Hz, 60 pulses at 20 Hz, 150 pulses at 50 Hz and 300 pulses at 100 Hz 1 ms-long pulse stimulations. This was done while maintaining the cells at -60mV in current-clamp mode. Current steps were injected in order to verify the identity of the cell.

##### For chemogenetic construct validation

action potentials were initiated via a series of depolarising current steps (from 0 to 200 pA increment 25 pA, 1 s long step) from a holding potential of – 60 mV.

##### Analysis

Analysis of action potentials was done using the event detection feature in Clampfit (Molecular Devices, v10.7) or custom Python scripts using a threshold detection above -20 mV. For the characterisation of IPSPs, maximum peak hyperpolarization was measured by subtracting the peak from the baseline of the response traces. The IPSP area was measured by summing the absolute value of each point subtracted by the baseline and dividing by the sampling frequency. The IPSP area was normalized to the peak hyperpolarization for each trace in order to compare different stimulation frequencies between cells.

### Immunohistochemistry

#### Immunofluorescence staining

NDNF-Cre mice aged 2-4 months, expressing either chemogenetic or optogenetic constructs were injected with a lethal dose of sodium pentobarbital (500 mg/kg) and transcardially perfused with cold 4 % paraformaldehyde (PFA) in phosphate buffered saline (PBS). Brains were removed, transferred to 4% PFA solution and stored overnight at 4°C. Using a vibrating microtome (Leica VT1000s), 70 to 100 μm thick horizontal or coronal brain slices were cut.

Free-floating sections were washed 3 x 5 minutes in 1 x PBS and incubated with a blocking solution (3 % goat serum, 0.3-0.5 % Triton X-100, 0.5 % bovine serum albumin, BSA, in PBS) at room temperature for 1 hour. Sections were then incubated with primary antibody in blocking solution (**Suppl. Table 2**) at 4°C overnight on a shaking platform. Primary antibody solutions were then removed, and sections were washed 3 x 10 minutes with PBS before incubation with a secondary antibody (**Suppl. Table 2**) for 3 hours at room temperature. Sections were then washed for a further 3 x 5 minutes in PBS then incubated with DAPI (Invitrogen, D1306 1:5000) for 5 minutes before a further 3 x 5-minute washes. Sections were then mounted onto glass slides using a hard set mounting medium (Fluoroshield, Sigma).

#### Image acquisition

Images were acquired on a Leica Mica or Leica LSM 170 confocal microscope through 20x (0.75 NA) non-immersive objective using laser wavelengths of 405 nm, 488nm and 594nm. The laser power and digital gain were adjusted for each staining condition to ensure optimal fluorophore visibility and kept the same within the same immunohistochemical condition.

#### Image analysis

The confocal images were processed using the FIJI software. After opening the images, the z-stack projections were collapsed into a single 2D image. To optimise image clarity and accuracy, the brightness/contrast was adjusted, background noise was subtracted. Neuronal quantification was performed manually using the “cell counter” analysis plugin.

### Statistical analysis

All statistical data analyses were performed using Prism, OriginPro software or Python scripts. Repeated two-way ANOVA, two-tailed unpaired and paired t-tests were used as appropriate. The paired and unpaired mean difference plots were generated using the Data Analysis with Bootstrapped ESTimation package (DABEST,^58^). 95% confidence intervals of the mean difference were calculated by bootstrap resampling with 5000 resamples. The confidence interval is bias-corrected and accelerated.

## Results

### NDNF-Cre targets layer 1 cells with a profile typical of NGF interneurons

We first sought to confirm that the NDNF-Cre driver line preferentially targets NGF cells using immunohistochemistry and electrophysiological characterisation following AAV-hSyn-FLEX-Chronos-GFP injection. Immunostaining of layer 1 of the visual cortex revealed co-expression of known molecular markers of NGF neurons, including reelin and NPY (**Suppl. Fig. 2A**) and little or no VIP co-expression. However, GFP expression was also seen in pyramidal shaped neurons within deeper layers (**Suppl. Fig. 2B**). In order to restrict gene expression to interneurons, we switched hSyn to the interneuron specific enhancer mDlx^59^. When AAV9-mDlx-FLEX-Chronos-GFP was injected in the superficial cortex of NDNF-Cre mice, GFP expression was almost exclusively seen in layer 1 (**Suppl. Fig. 2B**) with only occasional ectopic fluorescent cells (< 5 pyramidal shaped cells per spreading area). Layer 1 NDNF+ neurons exhibited either an early-spiking or a late-spiking phenotype when injected with depolarizing step current waveforms, as previously reported^18,19,23,60^. In addition, optogenetic depolarization of NDNF+ neurons evoked long-lasting inhibitory postsynaptic potentials (IPSPs) in pyramidal neurons, mediated by GABAB and GABAA_slow_ receptors (**Suppl. Fig. 2C, D**).

Taken together, the molecular and electrophysiological features of NDNF+ cells described here are consistent with those previously reported, and our intersectional strategy using NDNF marker in combination with an interneuron-specific enhancer, mDlx, allows us to selectively target NDNF+ interneurons in cortical layer 1.

### NDNF+ cell calcium fluorescence during epileptiform activity

While activity of PV+, SOM+, and VIP+ interneurons has been described during hyperexcitable states such as interictal discharges and seizures in ex vivo and in vivo animal models^28,29,33–35^, little is known about the activity of NDNF+ NGF cells in these conditions. We co-injected AAV9-hsyn-FLEX-GCaMP6f and AAV9-mDlx-FLEX-mCherry into the superficial layer of the visual cortex of NDNF-Cre mice in order to record calcium activity in NDNF+ neurons with concurrent electrocorticogram (ECoG) recordings. mCherry fluorescence allowed NDNF+ cells to be identified independently of their calcium activity. On the day of the experiment, pilocarpine was injected in layer 5 in the vicinity of the imaging window (∼1 mm away), to evoke focal epileptiform activity (**Fig. 1A**). As previously reported^37,61^, 20-30 minutes post-injection, interictal discharges emerged, which then evolved to frequent focal self-terminating seizures (**Fig. 1B, C**).

**Figure 1:**
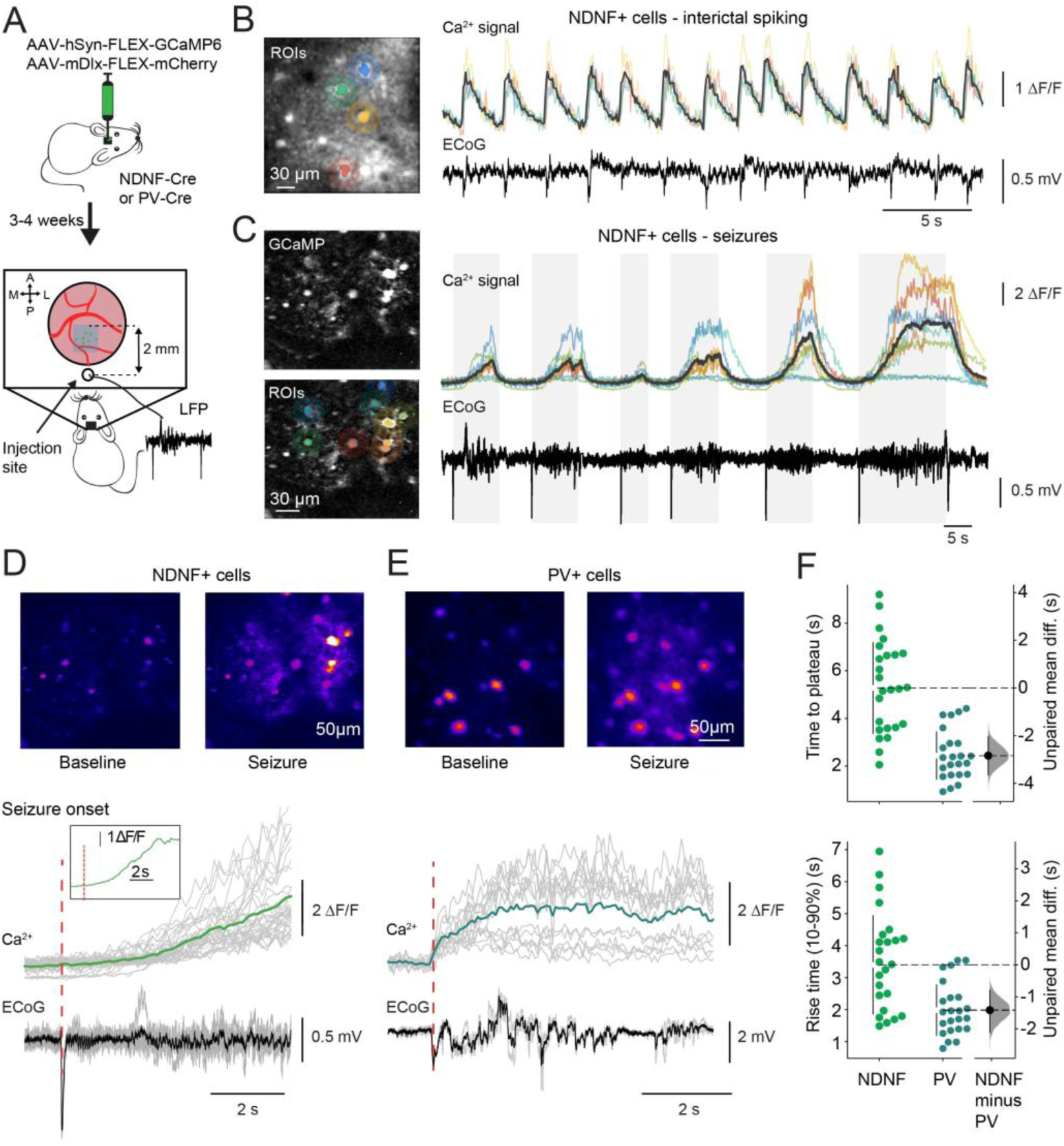
Delayed recruitment of NDNF+ neurons during seizures. (**A**) Experimental setup. (**B**) (Left panel) Representative field of view in the cranial window and multicoloured region of interest (ROI) masks on mCherry-positive neurons. (Right panel) Coloured lines depict calcium transients during interictal spikes from individual cells and correspond to neurons in the field of view (Left panel). The dark grey lines indicate average calcium transients for all neurons in the field of view. (**C**) (Left panel) Representative field of view (Top) and with ROI masks applied (Bottom). (Right panel) Calcium transients (Top) detected in NDNF+ neurons in response to pilocarpine-induced seizures recorded by ECoG (bottom). Coloured lines correspond to calcium transients from individual neurons depicted by coloured masks on image to the left. Dark grey line indicates average calcium fluorescence from all neurons within the field of view. (**D**) (Top panel) Calcium-dependent fluorescence during baseline (Left) and seizure episodes (Right) in an NDNF-Cre mouse. (Bottom panel) Average ECoG recording (black line) and calcium transient (dark green line) detected from neurons across multiple seizures within one imaging period (120 seconds). Individual seizures are shown in grey. (Inset) Average calcium response on an expanded time scale. (**E**) Calcium fluorescence in PV+ interneurons studied as in (D). Average calcium transient in PV+ cells shown in blue. (**F**) Average time for the calcium transient to plateau from seizure start in NDNF+ (green) and PV+ (blue) cells (Top panel) and average 10-90% calcium transient rise time (Bottom panel). The time to plateau unpaired mean difference is -2.84 s [95%CI, -3.65, -2.04, p<0.001, two-sided permutation t-test and p<0.001, Students’s t-test] and the mean time rise unpaired mean difference is -1.42 s [95%CI, -2.11, -0.79], p<0.001, two-sided permutation t-test, and p<0.001, Student’s t-test]; NDNF-Cre n=26 cells, 5 mice; PV-Cre n=26 cells, 3 mice.

Calcium transients were observed in NDNF+ cells during both interictal spiking (**Fig. 1B**) and seizures (**Fig. 1C**). We estimated the latency from seizure onset (defined by a sentinel spike) to the maximal calcium transient and compared between NDNF+ cells and PV+ neurons, which are rapidly recruited during seizures^28,33^. The recruitment latency of NDNF+ cells was more than twice that of PV+ neurons (**Fig. 1D-F**). The rise time was also longer in NDNF+ neurons than in PV+ cells (**Fig. 1G**), confirming their delayed recruitment during seizures.

NDNF+ NGF cells are thus recruited relatively slowly during focal seizures.

### Optogenetic hyperpolarization of NDNF+ cells during epileptiform activity

We asked if NDNF+ cells restrain or promote network activity during pilocarpine-induced interictal spiking and seizures by activating an inhibitory opsin in awake, head-fixed mice. The hyperpolarizing opsin ArchT was expressed in the visual cortex using a mDlx-Cre-dependent viral vector (**Fig. 2A**), and activated with 570 nm laser illumination delivered via an optic fibre placed above a cranial window. This ensured that the LFP recording and hyperpolarized NDNF+ cells were within the seizure focus. We separately verified that activation of ArchT resulted in hyperpolarization and suppression of spiking of NDNF+ cells using whole-cell patch clamp recordings in ex vivo brain slices (**Suppl. Fig. 3**).

**Figure 2:**
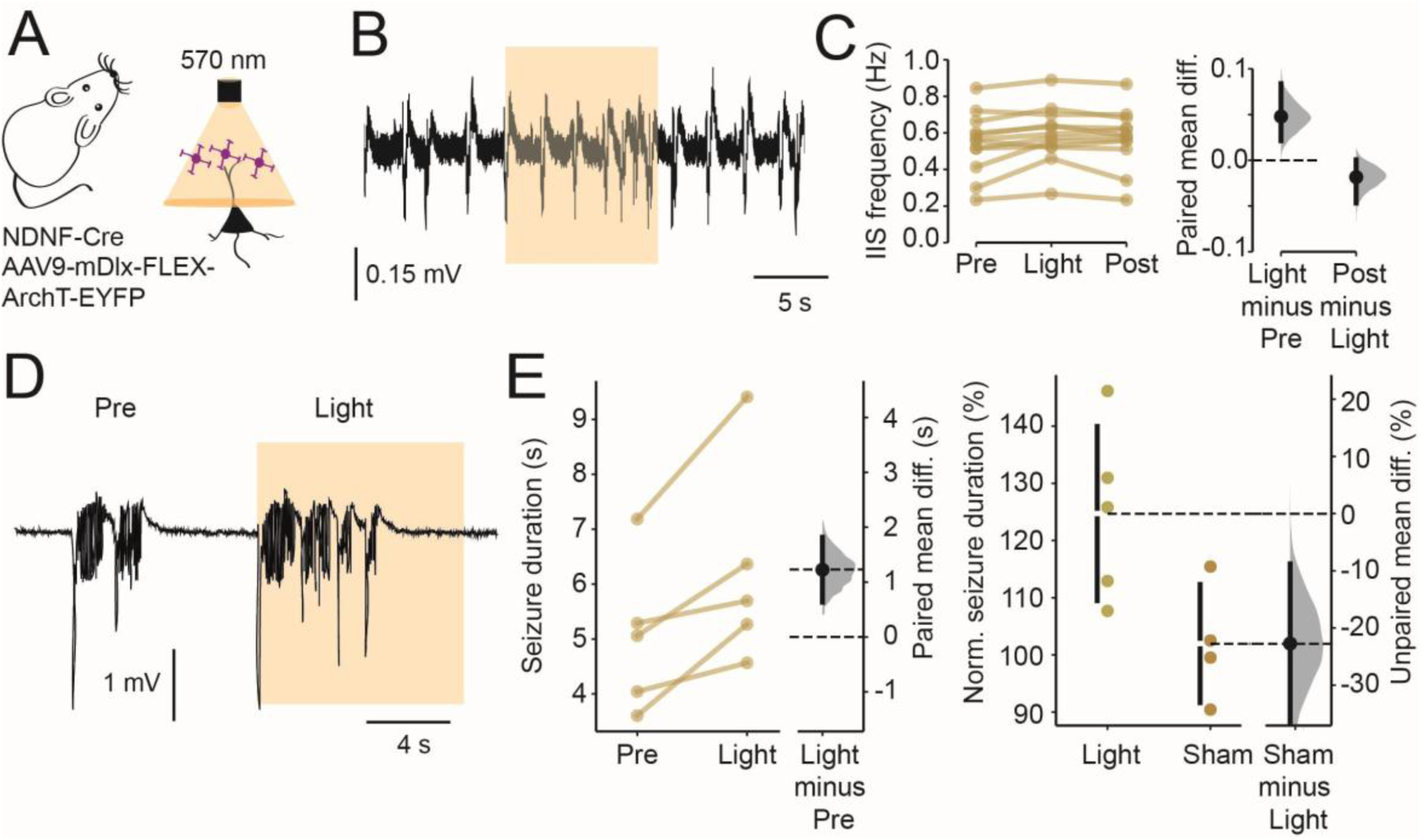
Optogenetic hyperpolarisation of NDNF+ neurons exacerbates epileptiform activity. (**A**) Experimental setup. (**B**) Representative ECoG showing interictal spiking evoked by pilocarpine injection. Yellow shading indicates a 10 s long 570 nm illumination of the cranial window. (**C**) Interictal spike (IIS) frequency increased during light exposure (the paired mean difference between pre-light and light period is 0.05 events [95%CI, 0.02, 0.08], p=0.009, two-sided permutation t-test, and p=0.013 paired t-test) and between light and post-light periods is -0.02 [95%CI, -0.05, 0.001], p=0.16, two-sided permutation t-test, and p= 0.165 paired t-test, n= 12 mice). (**D**) Representative seizures before and during NDNF+ cell optogenetic hyperpolarisation. Yellow shading indicates ‘light on’ period. (**E**) Seizure duration was increased during illumination compared to both a preceding seizure (Left panel, the paired mean difference between the duration of the preceding seizure and during NDNF+ cell activation is 1.22 s [95%CI, 0.63, 1.82], p=0.067, two-sided permutation t-test, and p=0.024 paired t-test) and sham laser activation (Right panel, the unpaired mean difference between the duration of sham and light stimulation is 22.73% [95%CI, 8.2, 38.2], p=0.0474, two-sided permutation t-test, and p=0.039 Student’s t-test, Sham n = 4 mice, Light n = 5 mice.)

Optogenetic hyperpolarization of NDNF+ cells *in vivo* resulted in a small but significant increase in the frequency of interictal spikes (**Fig. 2B, C**). This contrasts with the effect of optogenetic hyperpolarization of PV+ cells, which had very little effect on interictal spiking activity (**Suppl. Fig. 4**).

To investigate the effect of optogenetic hyperpolarization of NDNF+ cells during focal seizures, we triggered the laser upon detection of a sentinel spike in real-time. Consistent with the effect on interictal discharges, hyperpolarising NDNF+ cells resulted in an increase in seizure duration (**Fig. 2D-F**), as determined by comparing either to the immediately preceding seizure where laser illumination was not delivered, or to the effect of sham activation of the opsin where the seizure was detected but the laser was not activated. Overall, optogenetic NDNF+ cell hyperpolarization prolonged the seizure duration by ∼20%.

Taken together, these results suggest that NDNF+ neurons locally restrain pathological activity during both interictal discharges and seizure activity.

### Optogenetic depolarization of NDNF+ cells during epileptiform activity

Since inhibiting NDNF+ neurons promoted epileptiform activity, we hypothesized that depolarizing NDNF+ cells would have the opposite effect. To test this prediction, we used the same experimental setup but instead expressed the depolarizing opsin Chronos in NDNF+ cells (**Fig. 3A**). We used the same FGPA system to detect the seizure onset and delivered 470 nm laser illumination via the optic fibre to activate Chronos.

**Figure 3:**
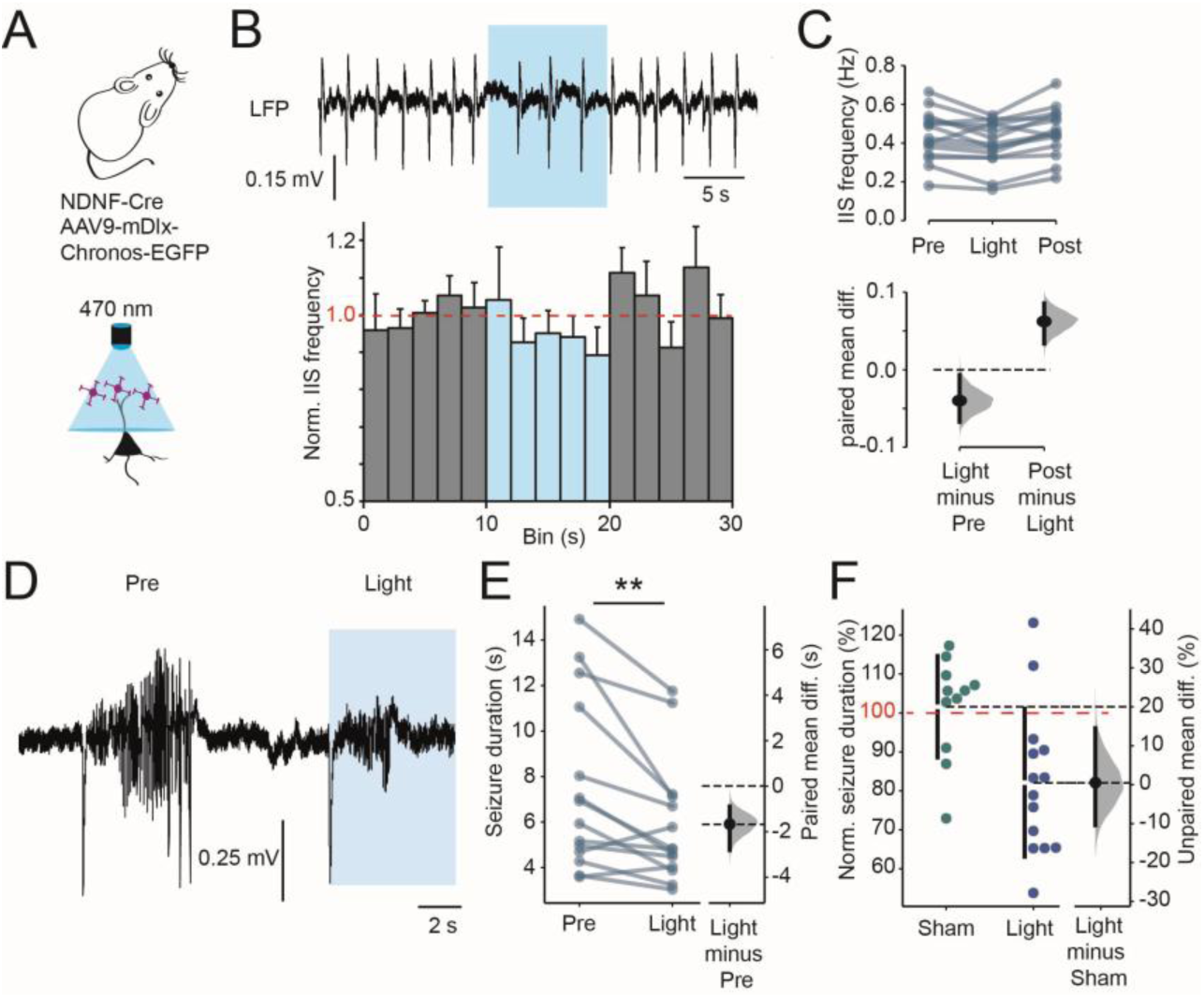
Optogenetic depolarisation of NDNF+ interneurons consistently reduces epileptiform activity. (**A**) Experimental setup. (**B**) Representative ECoG recording of interictal spiking following pilocarpine injection. Blue shading indicates 470 nm laser illumination via the cranial window. (**C**) Interictal spike (IIS) frequency decreased during activation of Chronos (Light) compared to a preceding or following 10 s period (the paired mean difference between pre-light and light period is -0.04 [95%CI, -0.07, -0.01], p=0.024 two-sided permutation t-test, and p=0.024 paired t-test; between light and post-light periods, the mean difference is 0.06 [95%CI, 0.03, 0.09], p=0.0014 two-sided permutation t-test, and p<0.001, paired t-test, between pre-light and post-light periods, the mean difference is 0.02 [95%CI, 0.01, 0.04], p=0.034 two-sided permutation t-test, and p=0.03, paired t-test n= 15 mice). (**D, E**) Seizure duration was decreased during Chronos activation compared to a preceding seizure (the paired mean difference between the duration of the preceding seizure and during NDNF+ cell activation is -1.66 s [95CI, -2.8, -0.9], p= 0.003 two-sided permutation t-test, and p=0.005 paired t-test, n=14 mice) or sham light activation (**F,** the unpaired mean difference between the duration of sham and light stimulation is -19.5% [95%CI, - 30.4, -5.50], p=0.009 two-sided permutation t-test, and p=0.008 Student’s t-test, Sham n=11 mice, Light n=14 mice).

We first evaluated several stimulus parameters to optimize the ability to activate NDNF+ cells optogenetically ex vivo. We confirmed that trains of 1 ms pulses elicited action potentials in NDNF+ cells using whole-cell patch clamp recordings. Most of the patched cells followed light pulses 1:1 up to 80-100 Hz (**Suppl. Fig. 5**). However, NGF cells are known to exhibit synaptic depression^26,62,63^. We therefore looked for a stimulating protocol that minimized depression of GABAergic synapses made by NDNF+ cells. Unexpectedly, when a 30 s gap was inserted between 2 stimuli, there was no detectable synaptic depression, and only when the gap was 10 s or less was depression observed. We also investigated whether high frequency stimulation sustained for several seconds resulted in strong depression.

While pulse trains elicited synaptic depression, the charge transfer was the biggest for 20 pulses delivered at 20-100 Hz (**Suppl. Fig. 5**). We thus used a refractory period of a minimum of 30 s between 2 stimulus trains to avoid depression of NDNF+ cell output. We also pseudo-randomly varied the stimulation frequency between 2, 4, 8, 10, 20, 50, and 100 Hz.

Overall, in vivo activation of NDNF+ cells using Chronos resulted in a consistent reduction in the number of interictal spikes (**Fig. 3B, C**) compared to the preceding 10 s period. The number of interictal spikes then returned to baseline levels in the 10 s period following cessation of light exposure. The effect was maximal at a stimulation frequency of 50-100 Hz (**Suppl. Fig. 6**). The effect of activating NDNF+ cells differed from that of activating PV+ or SOM+ cells because the latter resulted in an initial suppression of interictal spikes which faded and was quickly followed, especially for PV+ cells, by a large overshoot of activity upon terminating optogenetic depolarization (**Suppl. Fig. 7**). In contrast, NDNF+ cell stimulation continuously suppresses interictal spiking and only a small but significant rebound was observed after the light was switched off.

When optogenetic depolarization of NDNF+ cells was triggered by the sentinel spike, seizure duration was reduced by ∼20% when compared to the preceding seizure while sham stimulation had no effect (**Fig. 3D-F**). Consistent with the effect on interictal spiking activity, 50-100 Hz stimulation yielded the strongest effect (**Suppl. Fig. 6**).

Taken together, the optogenetic manipulations of NDNF+ cells had bidirectional effects on epileptiform activity in the visual cortex, consistent with a role in preventing runaway excitation.

### GABAB-receptor mediated persistent inhibition by NDNF+ cells during seizures

We previously showed that optogenetic activation of PV+ interneurons was anti-epileptic if delivered immediately at seizure onset but was paradoxically pro-epileptic if delayed by more than 2 s from seizure onset. This pro-epileptic effect was abolished when the potassium, chloride co-transporter 2 (KCC2) was overexpressed in pyramidal neurons, suggesting that the somatic transmembrane chloride gradient rapidly collapses during seizures, compromising the inhibitory effect of PV+ cells^37^. In the case of SOM+ interneurons, their anti-seizure effect also faded rapidly when optogenetic depolarization was delayed, although conversion to a pro-seizure effect was not observed. We speculated that, since NDNF+ neurons and NGF cells exert their inhibitory effects via dendritic GABAA_slow_ and GABAB receptors, their ability to inhibit pyramidal neurons may be less labile during seizures.

To test this hypothesis, we looked at the effect of optogenetic depolarization of NDNF+ cells delayed by >2 s from seizure onset, as we did previously for PV+ and SOM+ interneurons^37^. A robust ∼30% decrease in seizure duration was still obtained under these conditions (**Fig. 4 B, C**). Seizure duration did not change with sham optogenetic stimulation (**Fig. 4C**).

**Figure 4:**
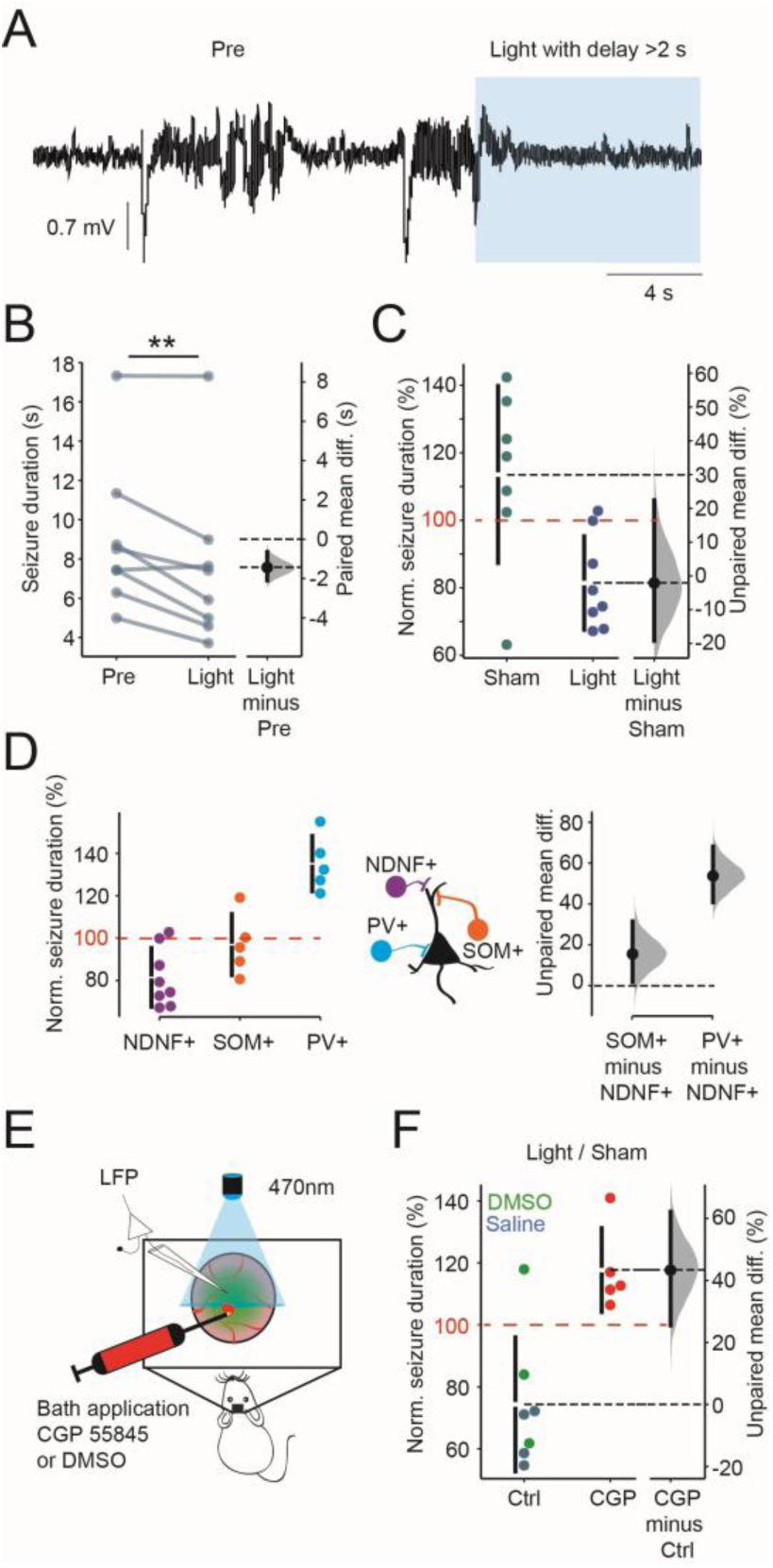
The anti-seizure effect of NDNF+ cells persists during seizures and depends on GABAB receptor activation. (**A**) Representative seizures recorded by ECoG from an NDNF-cre mouse. Blue shading indicates 470nm laser illumination to activate NDNF+ interneurons. (**B**) Seizure duration was significantly decreased by NDNF+ depolarization even when delayed by >2s from seizure onset (the paired mean difference between the duration of the preceding seizure and during NDNF+ cell activation is -1.44 s [95%CI, -2.09, -0.65], p=0.026 two-sided permutation t-test, and p=0.008 paired t-test, n=8 mice). (**C**) Normalised seizure duration was decreased during light exposure compared to sham light exposure (the unpaired mean difference between the duration of sham and light stimulation is -32.1% [95%CI, -49.25, -7.62], p=0.013 two-sided permutation t-test, and p=0.009 Student’s t-test, Sham n=7 mice, Light n=8 mice). (**D**) Normalised seizure duration following delayed optogenetic activation (>2s after seizure onset) of NDNF+, somatostatin+ (SOM+) or parvalbumin + (PV+) neurons. Delayed activation of NDNF+ neurons decreased seizure duration compared to both SOM+ and PV+ neurons (the unpaired mean difference of normalised duration between NDNF+ and SOM+ cell population is 15,49% [95%CI, 1.80, 31,39], p=0.075 two -sided permutation t-test, and p=0.079 Student’s t-test and between NDNF+ and PV+ cells is 53.70% [95%CI, 40.83, 68.12], p<0.001 tow-sided permutation t-test, and p<0.001 Student’s t-test, NDNF+ n=8 mice, SOM+ n=5 mice, PV+ n=5 mice; SOM+, and PV+ cell data from Magloire et al., 2019). (**E**) Experimental setup. Either the GABAB receptor blocker CGP 55845 (10 μM) or DMSO (0.1% in saline) control was applied topically via the cranial window. (**F**) Activation of NDNF+ neurons increased seizure duration in the presence of CGP 55845 (the unpaired mean difference between the control and CGP is 43.34% [95%CI, 25.5, 62.1], p= 0.009 two-sided permutation t-test, and p=0.003 Student’s t-test, Control n=7 mice, CGP n=5 mice).

NDNF+ interneurons thus retain their inhibitory action during seizure activity, in contrast to PV+ and SOM+ interneurons (**Fig 4D**, delay > 2s, data from ref. ^37^).

To assess whether this persistent anti-seizure effect of NDNF cells was mediated by GABAB receptors, we repeated the experiments whilst applying either the GABAB antagonist CGP55845 (CGP, 10 μM in 0.1% DMSO) or DMSO (0.1% in saline) or saline as control (**Fig. 4E**). To improve penetration of the drug, the dura was nicked at ∼5-6 sites. CGP consistently abolished the inhibitory action of NDNF+ neurons: activating NDNF+ cells in the presence of CGP resulted in an increase in seizure duration compared to the preceding seizure (**Fig. 4F**). Activation of NDNF+ neurons in the presence of DMSO continued to result in reduced seizure duration.

Taken together these results confirm that the inhibitory effect of NDNF+ cells on the local circuit persists even during excessive network activity such as ictal discharges, in contrast to PV+ and SOM+ cells, which represent about 70% of all interneurons^1^. This persistent inhibition is principally mediated by GABAB receptors which act independently of the postsynaptic chloride reversal potential.

### Chemogenetic activation of NDNF+ cells is anti-epileptic in ex vivo and in vivo recurrent seizure models

Optogenetics remains a vital tool to study the role of specific cells, but the requirement for an optic fibre and light source limits its translational potential. We therefore switched to a chemogenetic approach to activate NDNF+ neurons and interrogate its efficacy in preventing seizures. Chemogenetics not only permits titratable activation of neurons in response to a specific agonist but could also be used as an on-demand therapy, for instance to prevent impending status epilepticus^64^. We assessed the effect of chemogenetic NDNF+ neuron activation on seizure burden using both ex vivo slice models of seizures and an in vivo model of chronic epilepsy.

First, we verified that the excitatory DREADD hM3Dq can activate NDNF+ cells ex vivo. AAV9-mDlx-FLEX-hM3Dq-mCherry, or an mCherry control AAV, was injected into cortical layer 1 of NDNF-Cre mice, and after preparing acute brain slices whole-cell patch clamp was used to record action potential firing before and after the addition of the DREADD agonist clozapine-N-oxide (CNO, 10 μM). The resting input resistance and action potential firing were increased in neurons expressing hM3Dq-mCherry after the addition of CNO to the aCSF (**Suppl. Fig. 8**).

We next asked whether chemogenetic activation of layer 1 NDNF+ neurons reduces the duration of seizure-like events (SLEs) in a cortical brain slice model (**Fig. 5A**). Circuit excitability was increased by including a low concentration of the voltage-gated potassium channel blocker 4-aminopyridine (4-AP, 30 μM) in the aCSF, and SLEs were triggered by focal electrical stimulation in layer 5 of the cortex, while recording in an area of mCherry expression (**Fig. 5A**). We observed a non-significant reduction in seizure duration and in the number of spikes within a SLE, and a significant increase in interspike interval, compared to slices expressing the mCherry alone, when CNO was added to the aCSF (**Fig. 5B-E**).

**Figure 5:**
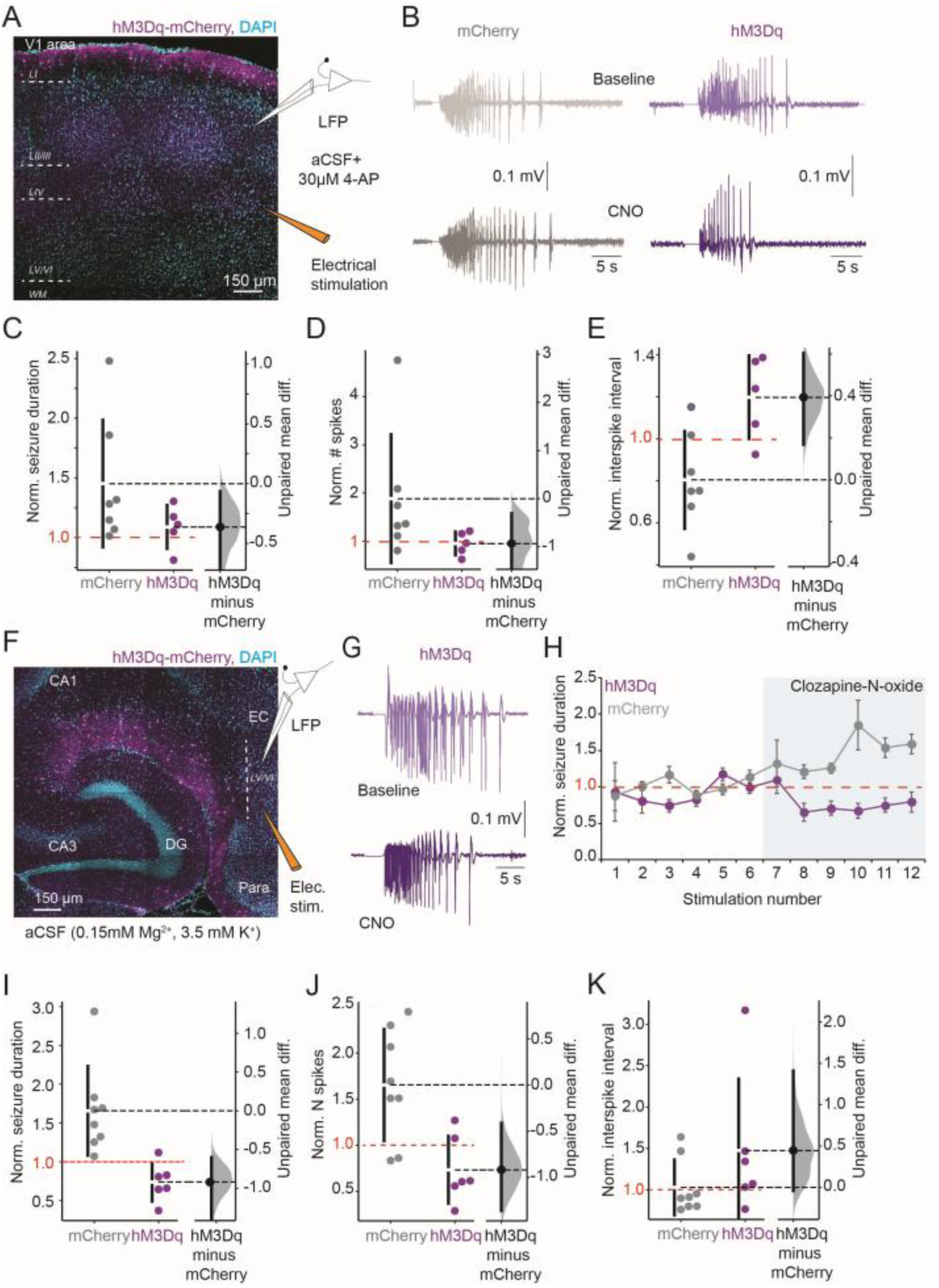
Chemogenetic activation of NDNF+ cells reduces seizure phenotype in an ex vivo model. (**A**) Cortical slice seizure model setup. (**B**) Representative LFP traces of seizure-like events (SLEs) evoked by electrical stimulation from slices expressing either hM3Dq-mCherry (purple), or mCherry only (grey) in NDNF+ neurons in baseline (Top panel) or following addition of 10 μM CNO (Bottom panel). (**C**) Seizure duration following 10 μM CNO application normalised to baseline (the unpaired mean difference between mCherry and hM3Dq groups is -0.36 [95%CI, -0.89, -0.06], p=0.19 two-sided permutation t-test, and p=0.18 Student’s t-test, mCherry n=7 slices; hM3Dq n=5 slices). (**D**) Normalised number of spikes per SLE baseline (the unpaired mean difference between mCherry and hM3Dq groups is -0.93 [95%CI, -2.44, -0.31], p=0.09 two-sided permutation t-test, and p=0.15 Student’s t-test, mCherry n=7 slices; hM3Dq n=5 slices). (**E**) Average normalised interspike interval per SLE after CNO application baseline (the unpaired mean difference between mCherry and hM3Dq groups is 0.39 [95%CI, 0.16, 0.60], p=0.01 two-sided permutation t-test, and p=0.01 Student’s t-test, mCherry n=7 slices; hM3Dq n=5 slices). (**F**) Hippocampal-entorhinal cortical slice seizure model. (**G**) Representative LFP recordings of stimulus-evoked SLEs from slices expressing hM3Dq before (top panel) and after (bottom panel) application of 10 μM CNO. (**H**) Duration of SLEs normalised to baseline (before CNO) in slices expressing mCherry only (grey; n = 8 slices) or hM3Dq-mCherry (purple; n = 6 slices). (**I**) Normalised seizure duration after CNO was decreased in hM3Dq expressing slices baseline (the unpaired mean difference between mCherry and hM3Dq groups is -0.92 [95%CI, -1.29, -0.60], p<0.001 two-sided permutation t-test, and p=0.003 Student’s t-test, mCherry n=8 slices; hM3Dq n=6 slices). (**J**) Number of spikes per SLE was decreased (the unpaired mean difference between mCherry and hM3Dq is -0.92 [95%CI, -1.36, -0.41], p=0.006 two-sided permutation t-test, and p=0.007 Student’s t-test mCherry n=8 slices; hM3Dq n=6 slices) and (**K**) interspike interval was increased in hM3Dq slices with CNO baseline (the unpaired mean difference between mCherry and hM3Dq is 0.44 [95%CI, -0.04, 1.41], p=0.19 two-sided permutation t-test, and p=0.20 Student’s t-test mCherry n=6 slices; hM3Dq n=6 slices).

GABAB receptor activation by baclofen is reported to be more potent in preventing epileptic activity in the hippocampus compared to neocortex^65^. Because NDNF+ NGF cell activation acts in large part via GABAB receptors, we speculated that a hippocampal seizure model would be more suited to test the efficacy of driving them chemogenetically. We previously used NDNF-Cre mice to target NDNF+ NGF cells in the stratum lacunosum moleculare (SLM, ref. ^60^, **Fig. 5F**) in the hippocampus, and found a similar electrophysiological profile as in the visual cortex. In addition, NDNF+ cells in the SLM strongly expressed the NGF cell marker reelin (**Suppl. Fig. 9**). Our intersectional approach therefore also works to target NDNF+ cells in the hippocampus. To test the impact of activating NDNF+ cells on hippocampal SLEs, we used low Mg^2+^ (0.15 mM) aCSF to increase neuronal excitability without initiating spontaneous seizures, and triggered SLEs with electrical stimulation of the entorhinal cortex. Following addition of CNO, the duration of SLEs and number of spikes within each SLE were decreased in slices expressing hM3Dq-mCherry, compared to slices expressing mCherry alone, which showed a slow increase (**Fig. 5G-K**). Chemogenetic activation of NDNF+ cells thus has an anti-seizure effect in both neocortical and hippocampal ex vivo models, and appears to be more robust in the latter.

Although NDNF+ cells are a potential cellular target for novel antiepileptic therapies, loss of interneurons in the hippocampus has been documented in temporal lobe epilepsy (TLE)^66–71^. Of the different populations, PV+ interneurons are the most vulnerable, although SOM+ cells are also affected (**Suppl. Fig. 10).** If NDNF+ NGF interneurons do not survive in chronic epilepsy, this may limit the effectiveness of a chemogenetic therapy targeting this cell type. We therefore asked whether NDNF+ cells are lost in a model of TLE evoked by intra-hippocampal injection of the chemoconvulsant kainate, which triggers a period of status epilepticus followed by frequent spontaneous focal and generalised seizures. The experiments were performed in mice where NDNF+ cells were labelled with mCherry as above. We measured the density of NDNF+ neurons in epileptic animals compared to intrahippocampal-saline injected control mice. We observed no reduction in the number of mCherry+ neurons (**Fig. 6A**) suggesting that NDNF+ NGF cells are relatively resistant to seizure-related damage in temporal lobe epilepsy.

**Figure 6:**
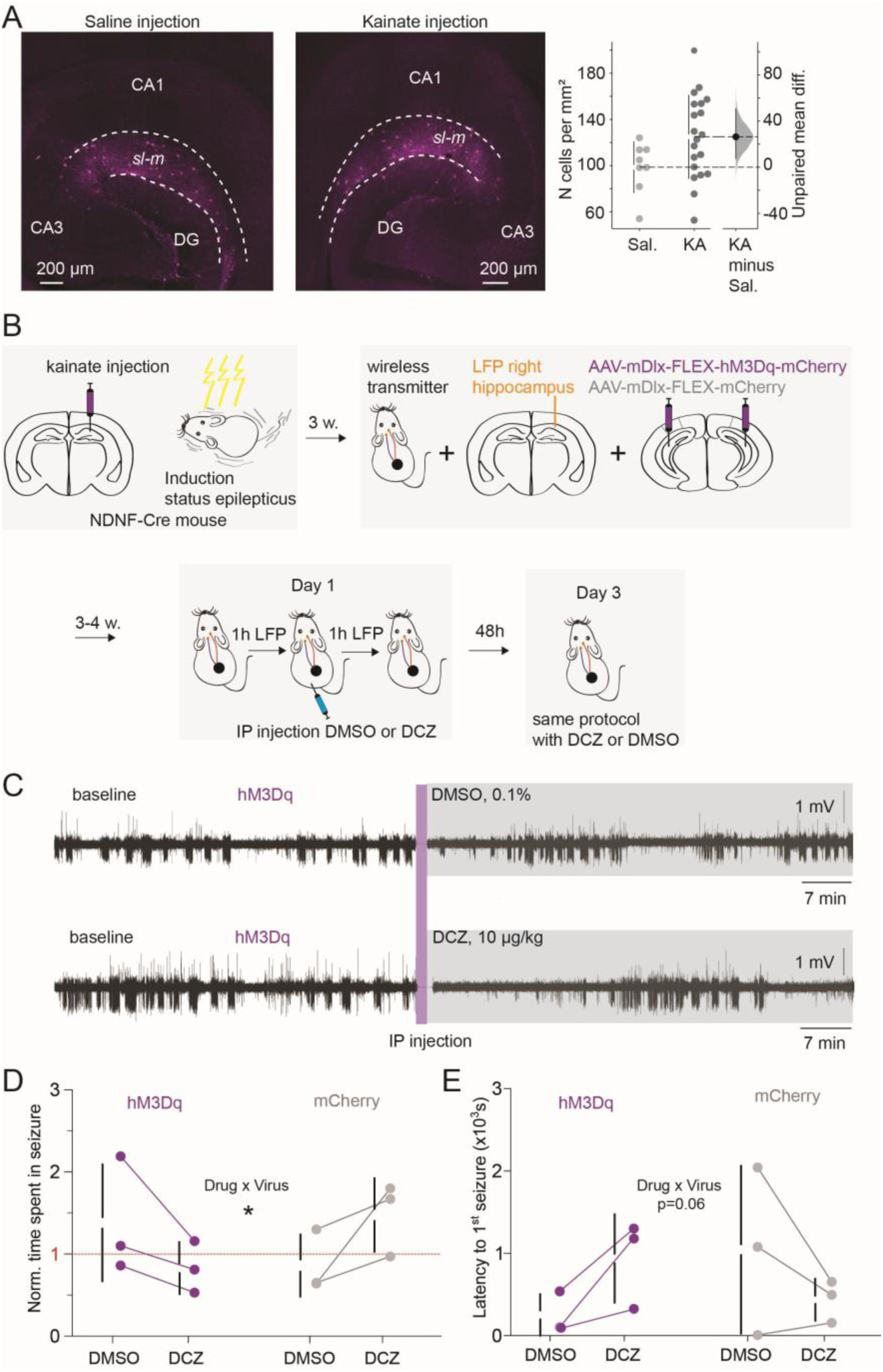
Chemogenetic activation of NDNF+ cells reduces seizure burden in an in vivo chronic model of temporal lobe epilepsy. (**A**) mCherry expression in NDNF+ neurons within the stratum lacunosum moleculare (sl-m) of the hippocampus from mice injected with either intrahippocampal saline (control) or intrahippocampal kainate (epileptic). DG = dentate gyrus. The number of mCherry-positive cells per area was unchanged in epileptic animals (the unpaired mean difference between Saline and Kainate is 26,3% [95%CI, 7.8, 50.5], p=0.072 two-sided permutation t-test, and p= 0.06 Student’s t-test, Saline n = 8 slices /3 mice; Kainate n= 20 slices /3 mice). (**B**) Experimental setup. (**C**) Representative LFP traces from hippocampal depth electrodes from a mouse expressing hM3Dq in NDNF+ neurons before and after intraperitoneal (i.p) injection of either DMSO (0.1% in saline) control or DCZ (10 μg/kg).

In a separate group of awake, head-fixed animals expressing both hM3Dq and GCaMP6f in cortical NDNF+ cells, we verified that the DREADD agonists clozapine (CLZ) and deschlorclozapine (DCZ) which activate hM3Dq in vivo directly, unlike CNO which undergoes back-conversion to clozapine^72^, increase the firing frequency of NDNF+ cells (**Suppl. Fig. 11**).

Finally, we asked whether chemogenetic activation of hippocampal NDNF+ cells could prevent spontaneous focal seizures in the TLE model. Three weeks following status epilepticus induction, we implanted epileptic animals with wireless transmitters connected to a hippocampal depth electrode to record focal epileptic activity, and injected either hM3Dq-mCherry or an mCherry-only virus bilaterally in the SLM of the hippocampus (**Fig. 6B**). Two weeks post-injection, the hippocampal LFP was recorded for one hour (baseline) and then again for an hour post-treatment with either DCZ (10 μg/kg in 0.1% DMSO, i.p.) or DMSO (0.1%) control. Two days later, the same recording was repeated with DMSO or DCZ switched, depending on what the mice received on the first day. The drug allocation was randomized and experimenters were blinded to both the viruses and drugs during the entire experiment and analysis. DCZ injection led to a decrease in seizure burden, defined as the time spent in seizure state, in mice expressing hM3Dq but not mCherry alone, compared to injection of DMSO (**Fig. 6C, D**), with a significant interaction between virus and substance injected (two-way ANOVA, **Fig. 6E**). The latency to the first seizure after drug injection was also increased, although this did not reach significance (**Fig. 6E**).

Activating NDNF+ interneurons chemogenetically thus has an anti-epileptic effect, identifying this population as a candidate cell target for diseases associated with circuit hyperactivity.

Vertical purple line represents time of injection and grey shading indicates post-injection ECoG recording. (**D**) Time spent in seizure was significantly decreased in mice expressing hM3Dq following injection of DCZ (two-way ANOVA, interaction p=0.033, n=3 mice per group). (**E**) Latency to 1^st^ seizure post-injection was increased in mice expressing hM3Dq following injection of DCZ (two-way ANOVA, interaction p=0.062, n=3 mice per group).

## Discussion

NDNF+ NGF cells have been identified as “master regulators” of the cortical column^9,26,73^, yet no study has previously examined their role in restraining excitation in the context of epileptiform hyperactivity. Using an intersectional interneuron-specific approach with an NDNF-Cre mouse line, we characterised the role of NDNF+ NGF cells in regulating cortical microcircuit excitability, both in extreme physiological and in pathological conditions. Through calcium imaging and optogenetic manipulation of NDNF+ cells, we demonstrated their substantial contribution to inhibitory restraint even several seconds after seizure onset, a mechanism mainly mediated by GABAB receptor activation. Finally, by enhancing NDNF+ cell intrinsic excitability using chemogenetics, we showed that this interneuron class continues to inhibit the cortical circuitry even in pathological epileptic conditions.

### NDNF molecular markers and NGF cell inhibition

We found that approximately 70% of NDNF+ cells express NPY, and 80% reelin. This compares with previous studies that report 30–70% of NDNF+ neurons expressing NPY, depending on the method used (e.g., NPY-GFP mouse line, RNA FISH^18,19,74^). In addition, a subset of NDNF+ cells exhibited a "late-spiking" (LS) phenotype, a hallmark of NGF cells^25^ whilst other cells displayed an "early-spiking" (ES) pattern. This dual spiking phenotype is consistent with other studies^17–19,26,74^. Indeed, Schuman et al. (2019) identified two types of NDNF-expressing cells, NGF and canopy cells, based on their co-expression of NPY, spiking patterns, connectivity profiles, and GABA signalling^19^. Despite the proposed distinction between these two cell types, their relative proportions within the NDNF+ population remain ambiguous. Our findings along with others, indicate that NPY and reelin are expressed in over 60% of all NDNF+ interneurons (80% for reelin;^18,26,74^), contrasting with the 30% reported by Schuman et al. (2019)^19^. Furthermore, Hartung et al. (2024), using unsupervised classification of NDNF+ neurons based on firing patterns and intrinsic properties, found no correspondence between NPY co-expression and the two electrophysiological cell types (ES vs. LS). They suggested that this discrepancy might be due to age-dependent NPY expression, with juvenile animals expressing less NPY^26^.

Although the precise proportion of NGF cells targeted by the NDNF-Cre mouse line remains unclear, canopy cells exhibit a lower connectivity rate with pyramidal neurons (18%) compared to NGF cells (72%), and generate smaller IPSCs that are mediated entirely by GABAA receptors^19^, suggesting a minor role in inhibiting the cortical column. The inhibitory output of the NDNF+ cell population in our study clearly evokes a powerful long-lasting inhibition mediated by both GABAA_slow_ and GABAB receptor activation, a signature unique to NGF cells^18,26,74^. Thus, while the NDNF+ cell population can be divided into at least two subtypes, it predominantly exhibits an inhibitory profile characteristic of NGF interneurons.

### NDNF+ interneuron recruitment during epileptic discharges

NDNF+ neurons are active during both interictal spikes and focal seizures originating approximately 1 mm away^61,75^, suggesting that, like PV+ and SOM+ cells, they can be recruited by local overexcitation^28,29,33–35^. However, their recruitment during focal seizures is significantly slower than that of PV+ cells. Several hypotheses could explain this slow-onset calcium rise in NDNF+ interneurons. Firstly, given that NDNF+ cells are located in layer 1 and receive afferent inputs from distal cortical and subcortical areas^18,74^, their gradual activation might result from the progressive recruitment of adjacent cortical territories as the seizure propagates. This mechanism alone is, however, unlikely to fully account for their slow-onset recruitment, as they are also active during local interictal events. A subset of NDNF+ NGF cells exhibit late-spiking activity^18,19,25,26,60,74^, which likely contributes to their delayed recruitment by slowly integrating changes in their membrane potential or incoming activity across hundreds of milliseconds to seconds. Secondly, NDNF+ cells potentially handle intracellular Ca²⁺ differently than do PV+ cells. This could result in slower Ca²⁺ entry and/or binding to GCaMP, thereby prolonging the rise time of somatic calcium signals independently of spiking activity. It is however unlikely that differences in intracellular calcium buffering account for this effect, since PV+ cells, despite their high expression of the calcium-binding protein parvalbumin, exhibit faster calcium signal rises. Lastly, NDNF+ NGF cells possess very short dendrites^17,21,25^ and demonstrate NMDA-dependent supralinear dendritic integration^76^. This suggests that NMDA receptor activation could serve as a potential source of Ca²⁺ entry, particularly during the spatiotemporally clustered synaptic inputs that would occur as seizures progress. Combined with their late-spiking profile and progressive recruitment from more distal regions, NMDA receptor activation may also contribute to the slow-onset activation of NDNF+ cells.

Our findings reveal a pattern of coordination where feedforward somatic inhibition by PV+ neurons would be followed by feedforward dendritic inhibition mediated by NDNF+ interneurons during epileptic activity, similar to previous observations between PV+ and SOM+ cells^29,34^.

### NDNF+ cell mediated inhibition stably restrains cortical overexcitation in contrast to PV+ and SOM+ cells

The effect of optogenetic manipulation of NDNF+ cells contrasts with that of PV+ and SOM+ cells. Indeed, optogenetic depolarisation of NDNF+ interneurons persistently suppressed both interictal spikes and seizures, irrespective of the timing of stimulation. While photo-depolarization of NDNF+, PV+, and SOM+ cells initially suppresses interictal spiking activity, their activation induces an important rebound of activity by the end of stimulation.

Consistently, while photo-depolarisation of all three interneuron classes at seizure onset exerts an anti-epileptic effect, their effects diverge markedly when depolarisation is delayed a few seconds into a seizure. NDNF+ cell activation elicits a persistent suppressive effect even when stimulated several seconds after seizure onset, whereas SOM+ cell excitation loses efficacy, and PV+ interneuron activation paradoxically exacerbates seizures.

Although the brain region, seizure model and optogenetic method were consistent across all experiments, several protocol differences must be acknowledged. PV+ and SOM+ cells are located in layers 2/3, 4, and 5, whereas NDNF+ cells are almost exclusively in layer 1^18,19,74^. This makes it challenging to achieve identical illumination configurations across experiments. Optogenetic manipulations of PV+ and SOM+ cells were thus performed with fibres inserted into layer 2/3, while NDNF+ cell experiments were conducted with the fibre positioned above the cortex. Additionally, optogenetic depolarisation of PV+ and SOM+ cells used continuous light illumination, whereas NDNF+ cells were stimulated intermittently. However, the inhibitory charge transfer elicited by continuous or discontinuous stimulation patterns of PV+ cells is equivalent^37^ and the greatest charge transfer for NDNF+ cell stimulation occurs at high frequency, which approximates continuous stimulation conditions. Thus, while minor differences exist in the optogenetic experimental protocols, they are unlikely to be the primary cause of the observed differences.

Our findings thus underscore the importance of the interneuron-specific spatiotemporal dynamics of inhibitory restraint in an hyperexcitable environment with NDNF+ cells being the most potent to maintain cortical excitability in check.

### Persistent NDNF+ cell anti-seizure effect is mainly mediated by GABAB receptor activation

In the present study, we show that NDNF+ cells persistently suppress seizure activity and that blocking GABAB receptors abolishes this anti-seizure effect. NDNF+ cells strongly inhibit not only pyramidal neurons but also PV+ cells (but not SOM+ cells)^18,26^.

Consequently, activation of NDNF+ cells would be expected to also hyperpolarize PV+ cells during seizures, contributing to an anti-seizure effect given the paradoxical role of this interneuron type^37,44^ particularly when applied more than 2 seconds after seizure onset^37^.

This phenomenon may contribute to the persistent anti-seizure effect observed here. However, when GABAB receptors are blocked, NDNF+ cell photo-depolarization promotes seizure activity in a manner qualitatively similar to that observed with PV+ cells. In our previous study, we demonstrated that somatic inhibition mediated by PV+ cells becomes pro-epileptic as seizures progress and that this switch was at least partially due to GABAA receptor mediated chloride loading of pyramidal cells^37^. Chloride loading renders GABAergic inhibition at best inefficient and at worst depolarising^77–79^. While chloride loading presents an attractive explanation for the pro-seizure effect of NDNF+ cells under conditions of GABAB blockade, this hypothesis remains to be investigated. Indeed, other possibilities cannot be discounted, such as chloride loading due to GABAA receptor activation leading to potassium efflux and depolarization of neighbouring neurons^80–82^.

Our findings thus strongly suggest that GABAB-mediated inhibition is critical for the suppressive role of NDNF+ interneurons under epileptic conditions.

### Implications for epilepsy

A relatively subtle increase in NDNF+ cell excitability by hM3Dq activation yielded a strong influence on cortical network excitability in both extreme physiological and pathological conditions. Although preliminary, these results suggest that enhancing the inhibitory function of NDNF+ cells could represent a viable therapeutic strategy for preventing seizures in TLE. Notably, NDNF+ neurons exhibit better survival in pathological conditions compared to PV+ and SOM+ cells^66–71^. Importantly, the NDNF marker is conserved in human NGF cells^83^, which could facilitate the translation of these findings into clinical applications. This is particularly relevant given the ongoing efforts in gene therapy for pharmaco-resistant epilepsy^84–86^.

Additionally, our data highlight the critical role of GABAB receptor activation in restraining hyperexcitation. While GABAB receptor agonists have shown deleterious effects in absence epilepsy^87^, accumulating evidence suggests their potential as therapeutic target in refractory partial epilepsy. GABAB agonists have demonstrated anti-seizure effects in various preclinical models of epilepsy^65,88,89^, and our findings provide further support for this approach.

Together, our results point towards NDNF+ cells and GABAB signalling as complementary and promising anti-seizure targets for the treatment of TLE.

## Acknowledgements

We are grateful to members of the Experimental Epilepsy Group at the UCL Queen Square Institute of Neurology for advice and technical assistance. This work was supported by Epilepsy Research UK, the Medical Research Council (MR/V034758/1, MR/W005204/1), the Wellcome Trust (212285/Z/18/Z), and the Rosetrees Trust. V.M. was the recipient of an Emerging Leader Fellowship from Epilepsy Research UK (F1901) and is the recipient of a Wellcome Career Development Award (227269/Z/23/Z). Y.S. received funding from the Japan Epilepsy Research Foundation.

