## Supplementary Data for "NDNF+ interneurons have a privileged role in regulating cortical excitability"

Supplementary tables and figures:

| Virus/ substance | Experiment | Coordinates | Volume per site | Dilution | Source |
|---|---|---|---|---|---|
| AAV9-hSyn-FLEX-GCaMP6f | Fig. 1 | **V1** AP -1.8 & -3.8 mm ML 2 mm DV -0.4 & -0.2 mm | 150 nL | 1:2 | Addgene 100833 (Stock titre: 2.2X1013 GC/mL) |
| | Supp. Fig.11 | **S1BF** AP 0 & -2 mm ML 3 mm DV -0.3 & -0.2 mm | 150 nL | 1:2 | |
| AAV9-mDlx-FLEX-Chronos-GFP | Fig. 3,4 and Supp. Fig. 1,2,5,6 | **V1** AP -1.8 & -3.8 mm ML 2 mm DV -0.4 & -0.2 mm | 150 nL | 1:2 | In house and VectorBuilder (Stock titre: $1X10^{13}$ GC/mL) |
| AAV9-mDlx-FLEX-ArchT-GFP | Fig. 2 and Supp. Fig. 3 | **V1** AP -1.8 & -3.8 mm ML 2 mm DV -0.4 & -0.2 mm | 150 nL | 1:4 | In house and VectorBuilder (Stock titre: $3.15x10^{13}$ GC/mL) |
| AAV9-mDlx-FLEX-mCherry | Supp. Fig. 8 | **V1** AP -1.8 & -3.8 mm ML 2 mm DV -0.4 & -0.2 mm | 150 nL | 1:2 | In house and VectorBuilder (Stock titre: $2.7x10^{13}$ GC/mL) |
| | Supp. Fig. 11 | **S1BF –** AP 0 & 2 mm ML 3 mm DV -0.3 & -0.2 mm | 150 nL | 1:2 | |
| | Fig. 5,6 and Supp. Fig. 9, 10 | **Hippocampus** AP -3.1 mm ML 3.12 mm DV -2.5 mm | 150 nL | 1:4 | |
| AAV9-mDlx-FLEX-hM3Dq-mCherry | Supp. Fig. 8 | **V1** AP -1.8 & -3.8 mm ML 2 mm DV -0.4 & -0.2 mm | 150 nL | 1:2 | In house and VectorBuilder (Stock titre: $1X10^{13}$ GC/mL) |
| | Supp. Fig. 11 | **S1BF** AP 0 & 2 mm ML 3 mm DV -0.3 & -0.2 mm | 150 nL | 1:2 | |
| | Fig. 5,6 and Supp. Fig. 9, 10 | **Hippocampus** AP -3.1 mm ML 3.12 mm DV -2.5 mm | 150 nL | 1:4 | |
| Kainic acid | Fig. 6 and Supp. Fig. 10 | **Dorsal hippocampus** AP -2.8 mm ML 3 mm DV -2 mm | Male: 140 nL Female: 80 nL | 1:3 | Tocris #7065 (Stock concentration 20 mM) |

***Table S1* – Viral constructs, kainate and injection sites** (V1: primary visual cortex, S1BF: primary somatosensory cortex, barrel field).

| Antibody | Species | Dilution | Blocking solution | Product code |
|---|---|---|---|---|
| Anti-parvalbumin | Mouse | 1:1000 | 0.5% Triton X-100<br>0.5% BSA in PBS | Sigma<br>P3088 |
| Anti-somatostatin | Rabbit | 1:200 | 0.5% Triton X-100.<br>0.5% BSA | Abcam<br>ab11912 |
| Anti-reelin | Mouse | 1:1000 | 0.2% Triton X-100.<br>0.5% BSA | Abcam<br>ab78540 |
| Anti-neuropeptide Y | Rabbit | 1:1000 | 0.3% Triton X-100.<br>0.3% BSA | Immunostar<br>22940 |
| Anti-vasoactive intestinal polypeptide | Rabbit | 1:500 | 0.3% Triton X-100.<br>0.3% BSA | Immunostar<br>20077 |
| Alexafluor488 anti-mouse | Goat | 1:1000 | PBS | Invitrogen<br>A11001 |
| Alexafluor594 anti-rabbit | Goat | 1:1000 | PBS | Invitrogen<br>A11012 |
| Alexafluor568 anti-rabbit | Donkey | 1:500 | PBS | Invitrogen<br>A10042 |
| Alexafluor568 anti-mouse | Goat | 1:500 | PBS | Invitrogen<br>A11004 |

***Table S2* – Antibodies**

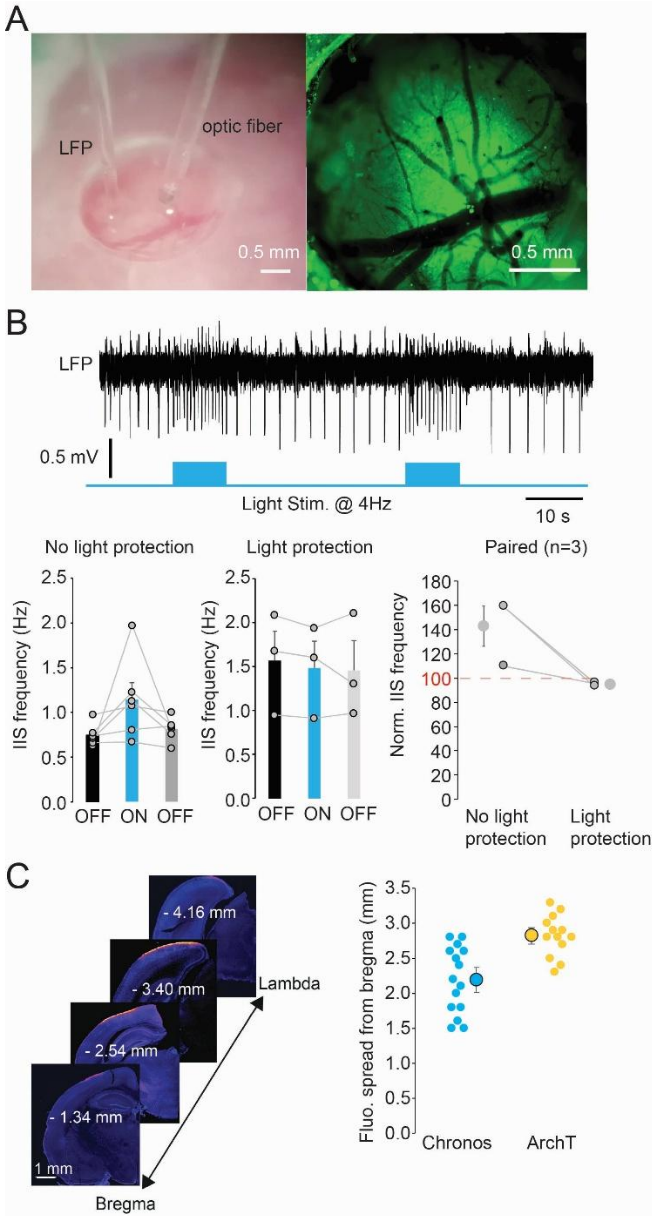


Supplementary Figure 1: **Experimental configuration for optogenetic manipulation of NDNF+ cells.** (**A**) (Left panel) Example cranial window performed in the superficial visual cortex for optogenetic experiments. (Right panel) Close-up photo of an optical window illuminated with 470 nm light. (**B**) (Top panel) Example LFP trace showing interictal spiking in the absence and presence of 10 s-long 470 nm light stimulation at 4 Hz (indicated by blue boxes). (Bottom panel) With no light shielding, laser illumination increased interictal spiking. This was prevented by shielding the eyes from the optic fibre light. (Right panel) Interictal spike frequency during light stimulation normalised to the 'light off' period.  (**C**) Images of successive brain sections from Bregma to Lamda taken from an NDNF-Cre mouse injected with AAV9-FLEX-mDlx-Chronos-GFP. Layer 1 NDNF+ neurons were positive for GFP (shown in orange). Chronos-GFP and ArchT-GFP expression had an anterior to posterior spread of 2.19 ± 0.12 mm and 2.82 ± 0.08 mm, respectively (average ± SEM). Chronos n = 15 mice; ArchT n = 13 mice.

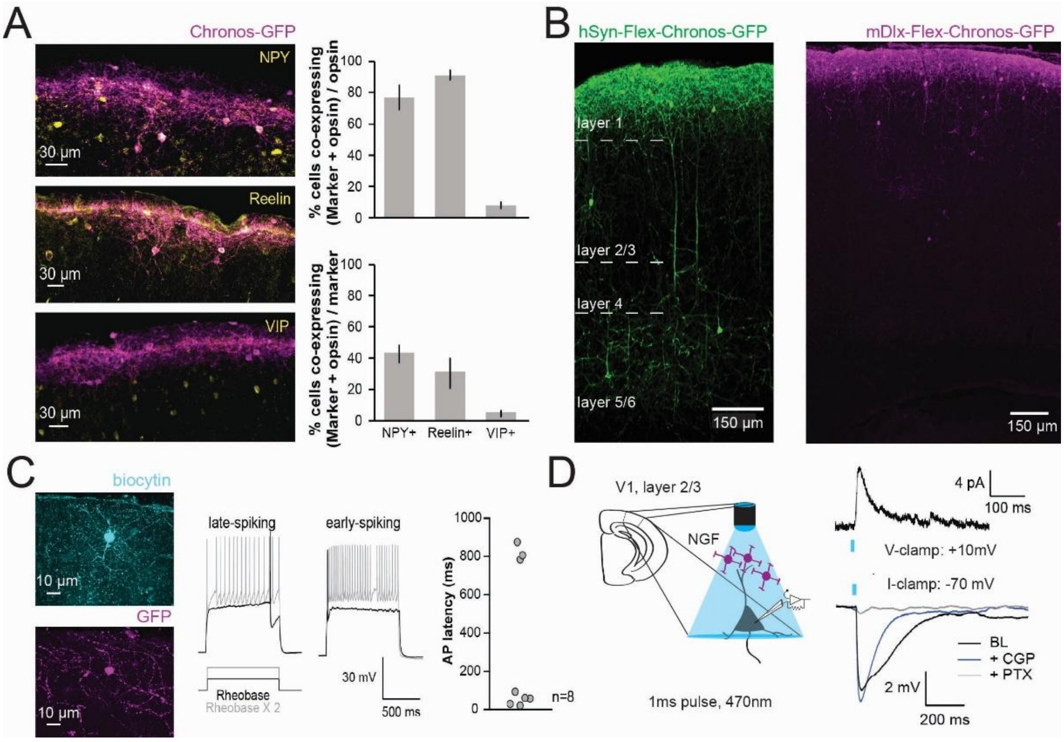


Supplementary Figure 2: **NDNF-Cre allows layer 1 NGF cells to be targeted.** (**A**) Co-staining of Chronos-GFP (pink) with known molecular markers of NGF neurons, neuropeptide Y (NPY; Top, yellow) and reelin (Middle, yellow) and with a marker not present in NGF neurons, vasoactive intestinal polypeptide (VIP; Bottom, yellow). (Right, Top panel) Percentage of Chronos-GFP positive neurons expressing each of the three markers. n = 3 slices, 2 mice. (Right, Bottom panel) Percentage of cells expressing each marker that also express Chronos-GFP. n = 3 slices, 2 mice (**B**) (Left) Example GFP expression (green) in the cortex of an NDNF-Cre mouse injected with an AAV containing a non-specific neuronal promoter, AAV9-hSyn-flex-Chronos-GFP. Note that some expression is seen within pyramidal shaped neurons in cortical layers 5/6. (Right). Example GFP expression (magenta) in the cortex of an NDNF-Cre mouse injected with an AAV containing an interneuron specific promoter, AAV9-mDlx-flex-Chronos-GFP. Expression is restricted to layer 1 interneurons. (**C**) A patched NDNF+ cell in cortical layer 1 expressing Chronos-GFP (magenta) labelled with biocytin (cyan) via the patch pipette. Patched cells showed two characteristic firing patterns, late-spiking and early-spiking. (**D**) Schematic of experimental setup. (Right, Top*)* Inhibitory post-synaptic current (IPSC) recorded in voltage clamp from a pyramidal neuron in response to activation of layer 1 NDNF+ neurons exhibiting a slow GABAA current. (Right, Bottom) Inhibitory post-synaptic potential (IPSP, black trace; BL - baseline) recorded in current clamp in response to activation of layer 1 NDNF+ neurons. Blocking GABAB receptors with CGP 55845 (navy) abolished the slow component of the IPSP. Addition of picrotoxin abolished the residual IPSP.

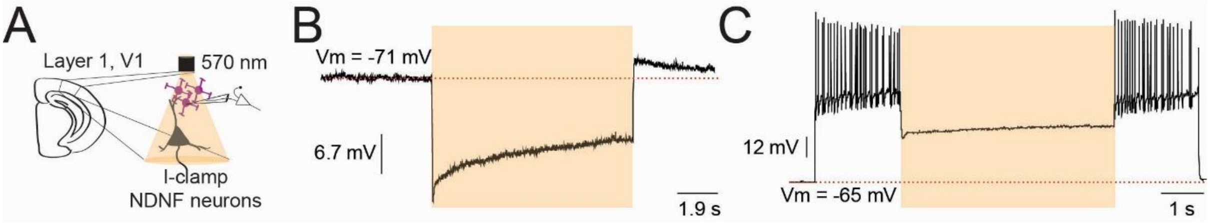


Supplementary Figure 3: **NDNF+ cell optogenetic hyperpolarisation with mDlx-FLEX-ArchT-GFP.** (**A**) Experimental setup. (**B**) Example of light-induced membrane hyperpolarisation recorded from an NDNF+ neuron expressing ArchT-GFP from a resting potential of -71 mV. Yellow shaded area 10s-long illumination. (**C**) Example of voltage trace from an NDNF+ neuron during injection of a depolarising current step to elicit action potentials. Laser activation (yellow shaded area) inhibited firing.

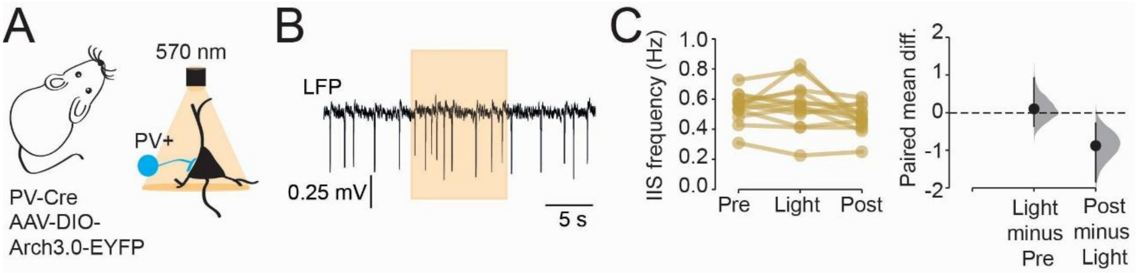


Supplementary Figure 4: **Effect of optogenetic hyperpolarization of PV + cells on interictal spiking.** (**A**) Experimental setup. (**B**) Example of LFP trace showing interictal spiking following pilocarpine injection. Illumination of PV neurons with 570 nm light (yellow shaded area, 10 s) led to a small increase in interictal spiking. (**C**) Overall, the interictal spike frequency across experiments did not change with light activation and exhibited a borderline-significant subsequent decrease (the paired mean difference between Pre and Light is 0.01 [95%CI, -0.04, 0.09], p = 0.79, two-sided permutation t-test, and p = 0.75, paired t-test; between Post and Light -0.09 [95%CI, -0.18, -0.03], p = 0.042 two-sided permutation t-test, and p = 0.045 paired t-test; n= 13 mice).

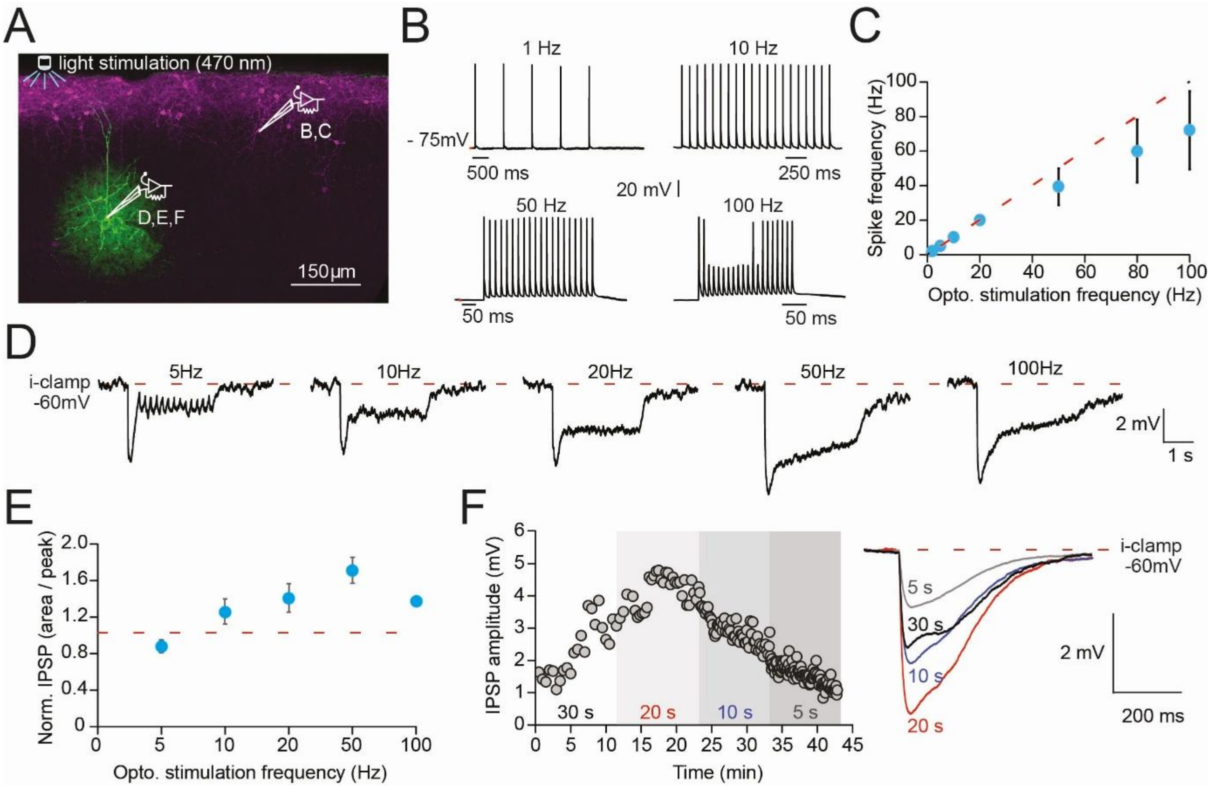


Supplementary Figure S5: **Optogenetic stimulation of NDNF+ cells: exploration of parameter space.** (**A**) Chronos-GFP expression in NDNF+ neurons in layer 1 of visual cortex (magenta) together with a patched pyramidal neuron loaded with biocytin (green). (**B**) Current clamp recordings from NDNF+ neuron expressing Chronos in response to 470 nm laser stimulation at different frequencies. (**C**) NDNF+ neuron firing frequency plotted against laser pulse frequency (n=4 cells). (**D**) Current clamp recordings from a neighbouring pyramidal neuron. (**E**) IPSP area normalised to peak hyperpolarization in response to optogenetic NDNF+ cell stimulation at different frequencies (n=7 cells). (**F**) Amplitude of IPSP in response to 1 ms light pulses every 30 seconds, 20 s, 10s or 5s.

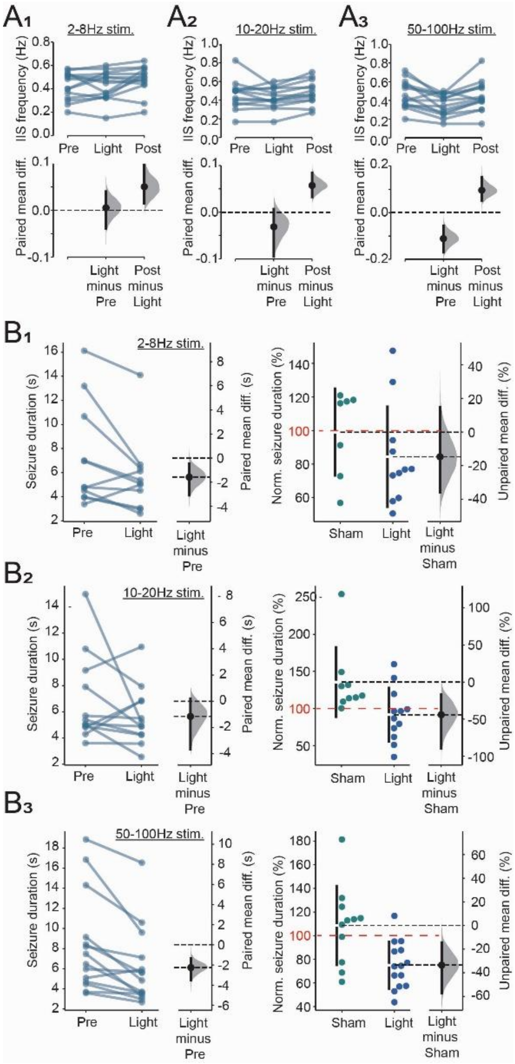


Supplementary Figure S6: **Frequency dependence of cortical network inhibition by optogenetic NDNF+ neuron depolarization.** (**A**) Interictal spiking following pilocarpine injection was unchanged in mice expressing Chronos-GFP in NDNF+ neurons when laser pulses were delivered at 2-8 Hz (paired mean difference between Pre and Light is 0.005 [95%CI, -0.038, 0.040, p=0.80 two-sided permutation t-test, and p= 0.79 paired t-test; mean difference between Post and Light is 0.05 [95%CI, 0.02, 0.10], p= 0.032 two-sided permutation t-test, and p=0.037 paired t-test, n=14 mice, $A_1$). When pulses were delivered at 10-20Hz, a non-significant decrease in spiking was observed, although the subsequent recovery reached significance (paired mean difference between Pre and Light is -0.03 [95%CI, -0.010, 0.007, p=0.28 two-sided permutation t-test, and p= 0.25 paired t-test; mean difference between Post and Light is 0.06 [95%CI, 0.03, 0.08], p= 0.003 two-sided permutation t-test, and p=0.004 paired t-test, n=13 mice, $A_2$). Optogenetic stimulation at 50-100 Hz yielded a significant decrease in interictal spike frequency (paired mean difference between Pre and Light is -0.11 [95%CI, -0.17, -0.06], p=0.003 two-sided permutation t-test, and p= 0.002 paired t-test; mean difference between Post and Light is 0.09 [95%CI, 0.05, 0.15], p= 0.002 two-sided permutation t-test, and p=0.003 paired t-test, n=14 mice, $A_3$). (**B**) Seizure duration exhibited a borderline-significant decrease when laser activation, triggered by the sentinel spike, was delivered at 2-8 Hz (the mean paired difference between Pre and Light is -1.58 s [95%CI, -3.08, -0.45], p=0.041 two-sided permutation t-test, and p=0.046 paired t-test, n=11 mice; Unpaired mean difference between Sham and Light is -14.7% [95%CI, -35.8, 14.6], p=0.29 two-sided permutation t-test, and p= 0.30 Student's t-test,

Sham n= 7 mice, Light n=11 mice, $B_1$). Similar results were obtained at 10-20Hz (the mean paired difference between Pre and Light is -1.16 s [95%CI, -3.63, 0.19], p=0.26 two-sided permutation t-test, and p=0.24 paired t-test, n=12 mice; Unpaired mean difference between Sham and Light is -44.0% [95%CI, -89.3, -16.8], p=0.013 two-sided permutation t-test, and p= 0.024 Student's t-test, Sham n= 9 mice, Light n=12 mice, $B_2$). Laser activation at 50-100Hz yielded a robust decrease in seizure duration (the mean paired difference between Pre and Light is -2.25 s [95%CI, -3.51, -1.34], p=0.001 two-sided permutation t-test, and p=0.002 paired t-test, n=14 mice; Unpaired mean difference between Sham and Light is -33.5% [95%CI, -57.3, -14.3], p=0.003 two-sided permutation t-test, and p= 0.005 Student's t-test, Sham n= 11 mice, Light n=14 mice, $B_3$).

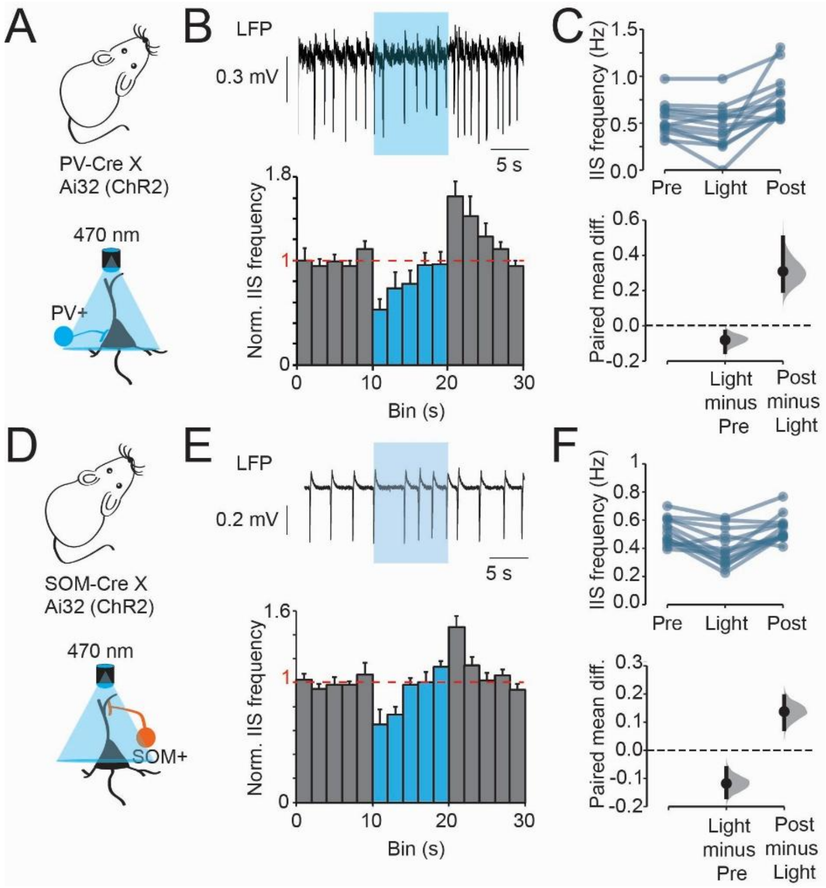


Supplementary Figure 7: **Effect of optogenetic depolarisation of PV+ and SOM+ cells on interictal spiking activity.** (**A**) Experimental setup for PV+ cell stimulation. (**B**) Representative LFP trace from a PV-Cre mouse showing interictal spiking following pilocarpine injection (Top panel). The blue box indicates time of light exposure. (Bottom panel) Normalised mean IIS frequency overall before, during and after light exposure (binning window 2s). (**C**) Interictal spiking was modestly decreased during light stimulation compared to a preceding 10 s period. Interictal spiking was increased following cessation of light stimulation compared to the illumination period. (Paired mean difference between Pre and Light is -0.08 Hz [95%CI, -0.15, -0.03], p= 0.012 two-sided permutation t-test, and p=0.021 paired t-test; paired mean difference between Post and Light is 0.31 [95%CI, 0.20, 0.50], p<0.001 two-sided permutation t-test, and p=0.002; paired t-test, paired mean difference between Pre and Post is 0.23 [95%CI, 0.15, 0.43], p<0.001 two-sided permutation t-test, and p=0.004 paired t-test, n=13 mice). (**D**) Experimental setup for SOM+ cell stimulation. (**E**) Example LFP trace from a SOM-Cre mouse (Top panel), (Bottom panel) Normalised mean IIS frequency overall before, during and after light exposure (binning window 2s) as in (B). (**F**) Interictal spiking was decreased during light stimulation (paired mean difference between Pre and Light is -0.12 Hz [95%CI, -0.17, -0.06], p= 0.002 two-sided permutation t-test, and p=0.001 paired t-test; paired mean difference between Post and Light is 0.14 [95%CI, 0.07, 0.19], p=0.002 two-sided permutation t-test, and p<0.001 paired t-test; paired t-test, paired mean difference between Pre and Post is 0.02 [95%CI, -0.004, 0.045], p=0.16 two-sided permutation t-test, and p=0.16 paired t-test, n=13 mice).

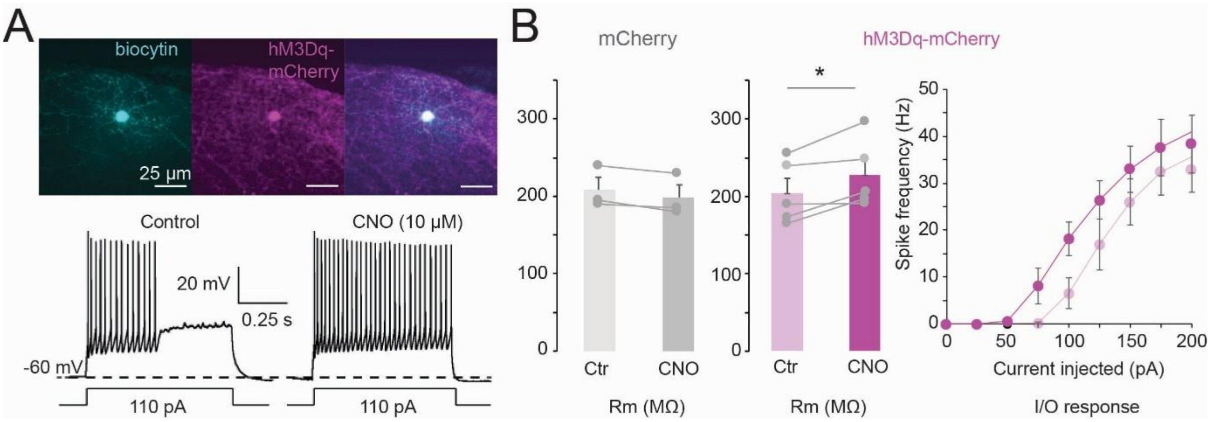


Supplementary Figure 8: **Activation of hM3Dq increases NDNF+ cell excitability ex vivo.** (**A**) Example NDNF+ neuron labelled with hM3Dq-mCherry (purple) that was patched and loaded with biocytin (cyan). (Bottom panel) Example current clamp recordings from the labelled neuron above. Action potentials elicited by depolarizing current injection before and after application of CNO. (**B**) Input resistance in NDNF+ neurons expressing either mCherry only (grey) or hM3Dq-mCherry (purple) before and after addition of CNO. CNO significantly increased the input resistance in neurons expressing hM3Dq (paired t-test, p=0.038, n=5 neurons). (Right panel*)* Spike frequency in response to increasing current step injections before (light purple) and after (dark purple) addition of CNO.

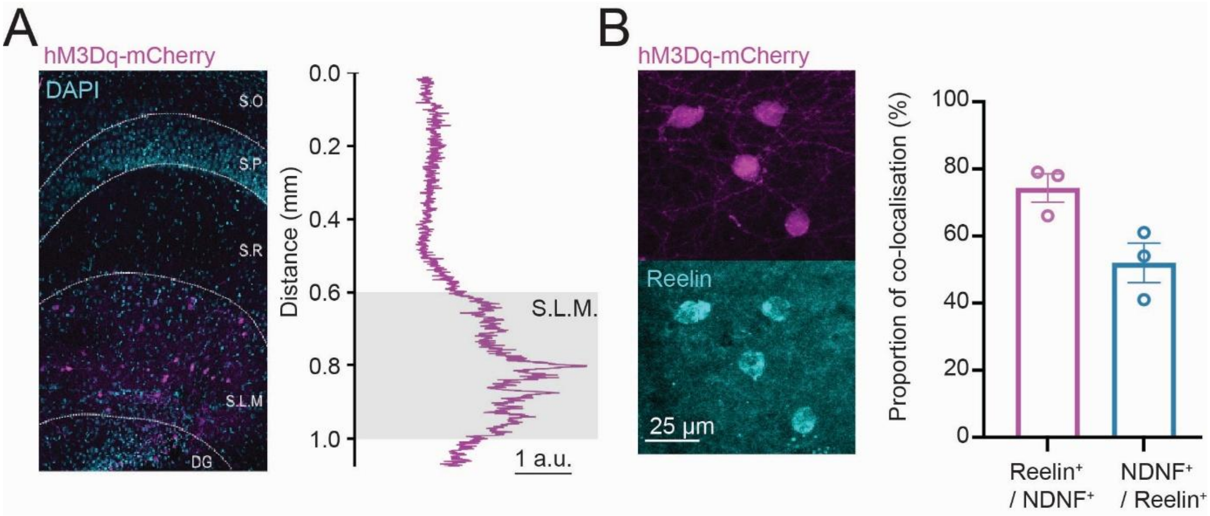


Supplementary Figure 9: **NDNF+ cell localisation and molecular profile in the hippocampus.** (**A**) mCherry expression (magenta) and DAPI staining (cyan) in the hippocampus of an NDNF-Cre mouse injected with AAV9-mDlx-FLEX-hM3Dq-mCherry. mCherry expression was restricted to the Stratum Lacunosum Moleculare (S.L.M) of the hippocampus. (Right panel*)* Fluorescence profile of mCherry staining throughout the layers of the hippocampus. Fluorescence was concentrated in the S.L.M. Abbreviations: S.O = Stratum Oriens, S.P = Stratum Pyramidale, S.R = Stratum Radiatum, DG = Dentate Gyrus. (**B**) mCherry expression (magenta) in NDNF+ neurons in the S.L.M co-stained with the NGF cell marker Reelin (cyan). The mean proportion of mCherry-positive cells expressing reelin was 74 ± 4.2% and the mean proportion of reelin cells expressing mCherry was 52 ± 5.5%, n = 3 mice 3 slices per animal.

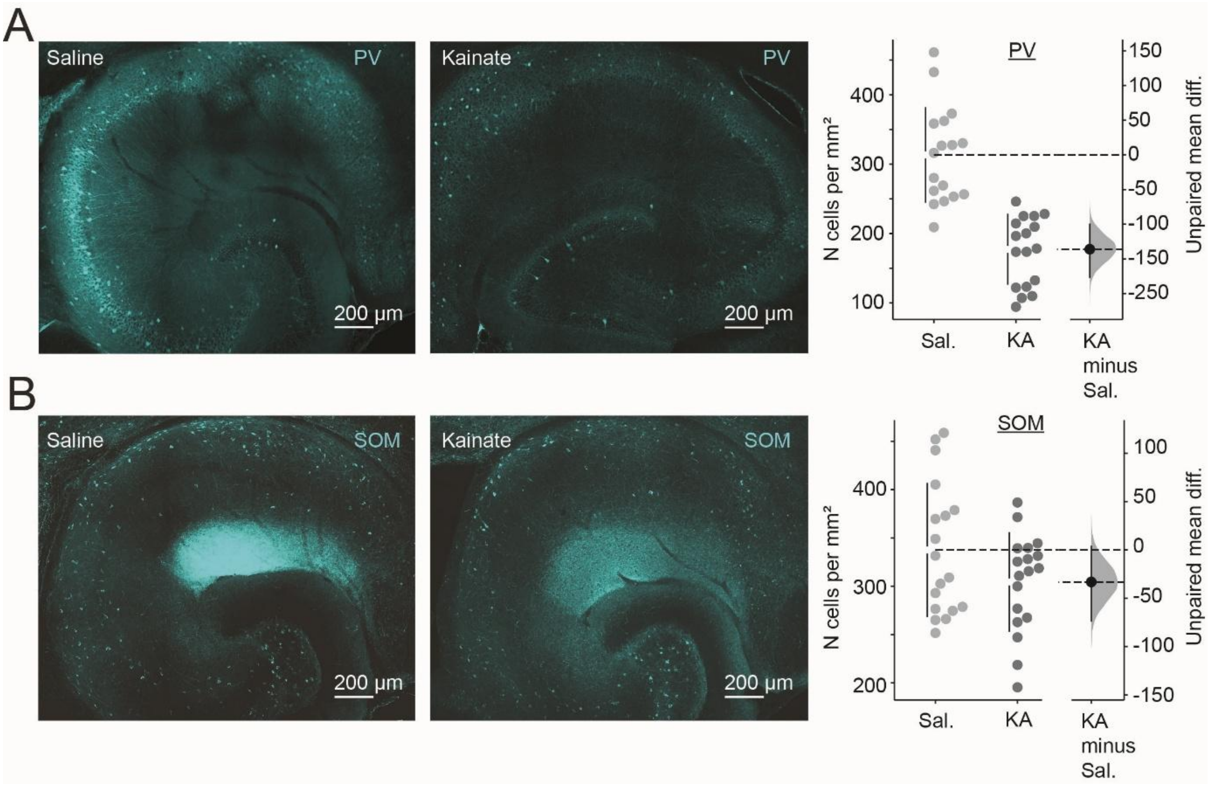


Supplementary Figure 10: **Loss of hippocampal interneurons in the intra-hippocampal kainate model of temporal lobe epilepsy.** (**A**) Representative images of parvalbumin (PV) staining in the hippocampus of a mouse injected with intrahippocampal saline (left) or kainate (right). The number of PV+ neurons was reduced in epileptic mice injected with intrahippocampal kainate (n=18 slices / 3 animals) compared to non-epileptic controls (n= 18 slices/ 3 animals; unpaired mean difference is -136 cells [95%CI, -176, -100], p<0.001 two-sided permutation t-test, and p<0.001 Student's t-test,). (**B**) Somatostatin (SOM) staining in the hippocampus of a non-epileptic saline injected control (left) or a kainate-injected epileptic mouse (right). The number of SOM+ interneurons was slightly reduced in epileptic mice (n = 18 slices / 3 mice) compared to saline-injected controls (n = 18 slices / 3 mice, unpaired mean difference is -33 cells [95%CI, -73, 4], p=0.11 two-sided permutation t-test, and p=0.11 Student's t-test).

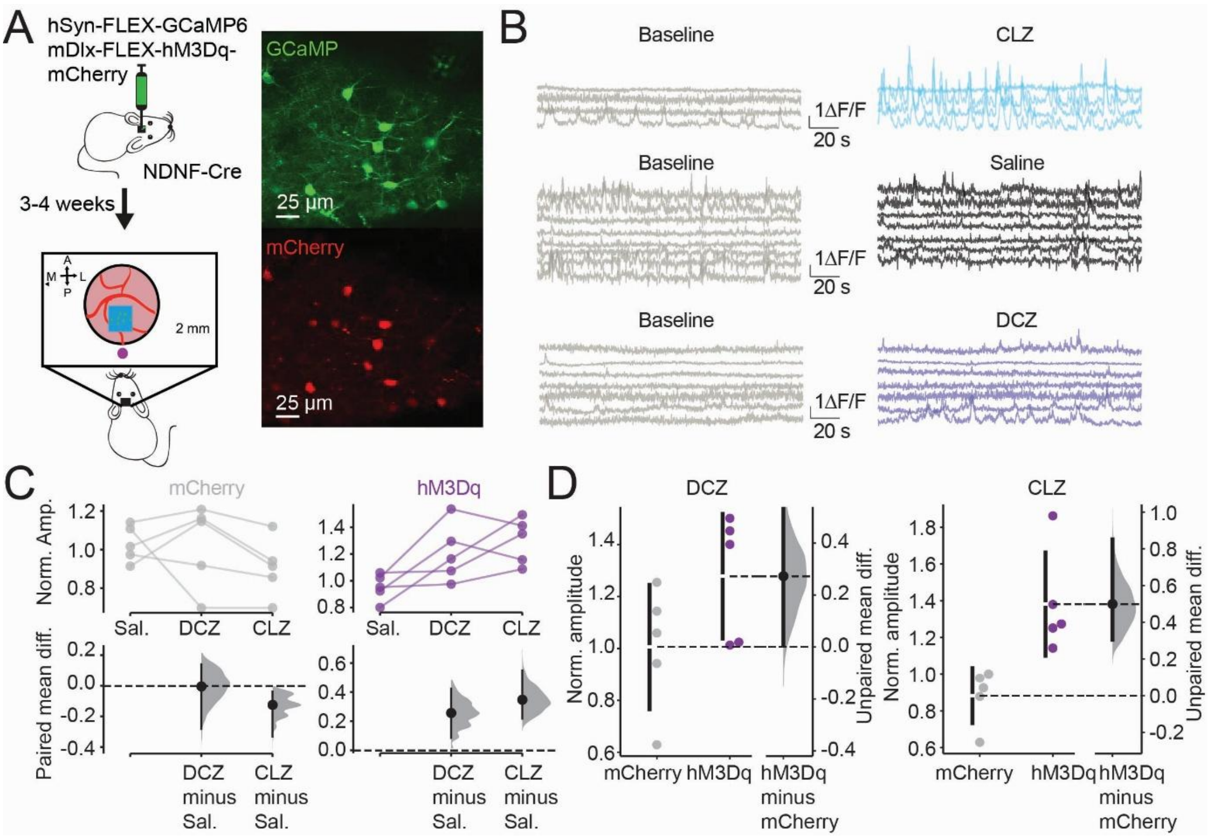


Supplementary Figure 11: **In vivo chemogenetic activation of NDNF+ cells.** (**A**) Schematic of experimental setup. (Right panel) Example GCaMP6f expression (green) and mCherry expression (red) in NDNF+ neurons within a field of view. (**B**) Representative calcium traces from a single mouse before (baseline) and after injection of clozapine (CLZ; blue), saline (dark grey) and deschlorclozapine (DCZ; purple). (**C**) Average amplitude of calcium transients following injection of DCZ, saline or CLZ normalised to the baseline period. In mice expressing mCherry only (grey) no change in amplitude was observed in the presence of DCZ or CLZ compared to saline (paired mean difference between Saline and DCZ is 0.004 [95%CI, -0.15, 0.24], p= 0.81 two-sided permutation t-test, and p= 0.97 paired t-test; paired mean difference between CLZ and Saline is -0.12 [95%CI, -0.33, -0.04], p<0.001 two-sided permutation t-test, and p=0.17 paired t-test, n= 5 mice). In animals expressing hM3Dq-mCherry, an increase in amplitude of calcium transients was seen with DCZ and CLZ compared to saline (paired mean difference between Saline and DCZ is -0.25 [95%CI, -0.42, 0.09], p<0.001 two-sided permutation t-test, and p= 0.06 paired t-test; paired mean difference between CLZ and Saline is 0.34 [95%CI, 0.21, 0.55], p=0.06 two-sided permutation t-test, and p=0.02 paired t-test, n= 5 mice). (**D**) Amplitude of calcium transients following injection of DCZ (left, unpaired mean difference between mCherry and hM3Dq is 0.27 [95%CI, 0.01, 0.55], p=0.10 two-sided permutation t-test, and p=0.11 Student’s t-test, n=5 mice per group) and CLZ (right, unpaired mean difference between mCherry and hM3Dq is 0.49 [95%CI, 0.30, 0.85], p<0.001 two-sided permutation t-test, and p=0.008 Student’s t-test, n=5 mice per group) normalised to baseline.
